# Suppressing the suppressor: Gallic acid induced asymmetric tetramerization of the pleotropic virulence factor SuhB from *Pseudomonas aeruginosa* abolishes its extragenic suppressor activities. A structure-based functional study

**DOI:** 10.1101/2025.11.12.687946

**Authors:** Vinay K. Yadav, Abinash Jena, Mitali Mukerji, Sudipta Bhattacharyya

**Affiliations:** Department of Bioscience and Bioengineering, Indian Institute of Technology, Jodhpur, Rajasthan (342037), India

## Abstract

*Pseudomonas aeruginosa* SuhB (PaSuhB) is a member of the bacterial Inositol monophosphatase family proteins. Numerous scientific evidences suggest PaSuhB is the pleotropic regulator of different metabolic pathways involved in bacterial biofilm formation and virulence determination. In this study, we have solved the high-resolution crystal structures of PaSuhB in its apo and substrate (Inositol monophosphatase) bound forms. Moreover, we carried out 3D pharmacophore modelling of the bound substrate to identify gallic acid, a phyto-phenol, abundant in medicinal plants, as a novel PaSuhB inhibitor. The high-resolution crystal structure of gallic acid/PaSuhB binary complex leads to the identification of a novel allosteric ligand binding site of the protein. *In vitro*, gallic acid induces the cold-sensitive growth of *P. aeruginosa* and *E. coli*, the previously reported phenomenon observed in *suh*B deletion mutants and also inhibits the swimming motility of *P. aeruginosa*. The plausible anti-bacterial molecular mechanism of action of gallic acid is presented herein.

---

*Pseudomonas aeruginosa* is a ubiquitous, encapsulated, Gram-negative, aerobic, rod-shaped bacterium known for its pathogenic potential in both plants and animals including humans. It is the member of the ESKAPE group which has been emerging as drug resistant pathogens at alarming rates throughout the globe. It majorly infects immunocompromised individuals and considered as one of the leading pathogens in hospital acquired infections [1-2]. The infections emerge due to *P. aeruginosa* that have been taken much attentions are chronic infections in cystic fibrosis (CF) patients and acute infections in hospitalized patients admitted due to burn injury. [3-4]. *P. aeruginosa* infections become deadliest due to emergence of highly drug resistant and virulent strains since it has inherent potential to form biofilm [5-6]. To cater these resistance *P. aeruginosa* infections there is no treatment options hence it is necessary and global demand in health sectors to find new potential drug targets and develop new small molecules-based inhibitors that can cope up this deadliest pathogen [7]. In bunch of the literatures, it is reported that inositol metabolic pathway regulates cell signalling, numerous physiological and pathophysiological activities in all form of lives [8-12]. Inositol monophosphatases (IMPases) are magnesium activated and lithium sensitive enzymes of FIG (Fructose 1,6 bisphosphatase, Inositol monophosphatase, GlpX) superfamily that hydrolyse inositol, sugar and nucleotide phosphates with promiscuous substrate specificity [13-16]. In eukaryotes, these enzymes are involved in phosphatidylinositol signalling and responsible for several physiological and pathophysiological conditions including bipolar disorders which represents major drug target for lithium therapy [17].

In prokaryotes, recent studies have found that IMPases are involved in several pathways which confers bacterial virulence like regulation of biofilm formation in Burkholderia and *Staphylococcus aureus*, regulation of exopolysaccharide biosynthesis in Rhizobium and *Staphylococcus aureus* [18-20]. It is also suggested that this enzyme’s orthologues are also involved in the mycothiol biosynthesis pathway which is major oxidative defence system of mycobacterium [21]. In *E. coli* this enzyme confers cold sensitivity and acts as extragenic suppressor of temperature sensitive phenotypes. Moreover, they play crucial roles in 30S ribosome biogenesis, RNA folding and maturation, and rRNA transcription anti-termination via interaction with NusA AR2 domain and weak interaction with RNA polymerase [22-24]. In Gram-negative bacteria myo-inositol and inositol metabolic pathways are limited which indicates that the enzymatic activity of IMPases are not the primary functions instead its regulatory functions are vital for bacterial virulence and drug resistance. In *Pseudomonas aeruginosa* IMPase (Gene ID: PA3818) are characterized as one of the well-known virulence factors with pleotropic regulatory functions. It is one of the global gene regulators and extragenic suppressor (PaSuhB) protein of temperature sensitive mutations. PaSuhB mutant analysis along with transcriptomic data suggest that this gene involves in regulation of bacterial cytotoxicity to human cells mediated through expression of type-III secretion system during acute infections of *P. aeruginosa*. Moreover, this protein plays crucial role in *P. aeruginosa* swimming motility and production of pyocyanin one of the most dangerous virulent factors which hijacks host defense system and promotes cytotoxicity during infections. All these aforementioned functions are attributed through GacS/GacA-RsmA pathway. Additionally, it also regulates biofilm formation during chronic infections like in cystic fibrosis patients through regulation of type-VI secretion system. Recently, one of the most critical roles of this enzyme is reported that it regulates the motile sessile switch in *Pseudomonas aeruginosa* through diguanylate cyclase and phosphodiesterase expression mediated via c-di-GMP pathway [25, 26]. In other study, it is evident that PaSuhB interact with ribosomes during protein translation and regulates expression of drug efflux pump MexXY which subsequently responsible for drug resistance to aminoglycoside and fluoroquinolones inversely increased drug sensitivity to beta-lactam antibiotics [27]. Despite the abundant pathophysiological regulatory roles and considerably attractive drug target the structure based functional characterization and small molecule-based inhibitors of PaSuhB is not reported yet. In the present paper we have solved high-resolution crystal structure of PaSuhB in its apo form and with its substrate D-*myo*-inositol 1-monophosphate (IPD) bound holo-structure. The IPD bound structure enabled us to conduct pharmacophore modelling of the bound substrate and led us to identify small molecules that may interact at the active site’s catalytic cleft of PaSuhB molecule. In this context, we identified gallic acid (GDE) a polyphenol highly abundant in medicinal plants and herbs that binds at the active site catalytic cleft of PaSuhB. GDE (3, 4, 5-trihydroxybenzoic acid) is a bioactive naturally occurring product and a generally recognized as safe compound to human use [28-29]. It is found in several medicinal plants that are used extensively in Ayurveda such as *Terminalia bellirica, Terminalia chibula, Phyllanthus emblica* and *Andrographis paniculata* etc [30-31]. GDE is known for its antibacterial, antiviral, antifungal, anti-inflammatory and anti-oxidative activities during recent decades, so it has a wide range of applications in food, medicine, biological and chemical industries [32-34]. Despite of tremendous therapeutic benefits exact molecular mechanism of GDE is still the challenge for scientific community. In the present study, we employed detailed structure based functional assays to uncover the plausible antibacterial mechanism of action of GDE. Our comprehensive findings corroborate GDE as the bonafide PaSuhB inhibitor and may serve as a potential lead scaffold to develop small molecule based therapeutic leads against MDR *P. aeruginosa* infections in the future.

## Results

### *P. aeruginosa* SuhB is structurally similar to typical IMPases with three metal binding sites at active site catalytic cleft

The earlier reported pleotropic virulence regulatory roles of PaSuhB in *P. aeruginosa* pathogenesis demand identification of novel small molecule-based inhibitors to cease the catalytic as well as virulence regulatory activities of PaSuhB. Intriguingly, pharmacophore guided inhibitor designing is one of the fascinating approaches which have high success rates to develop/design precise small molecule-based inhibitor leads. Hence, in this context, crystal structure of PaSuhB complexed with its substrate and metal ions may serve as the pre-requisite for pharmacophore guided ligand scouting. For this purpose, the ORF encoding PaSuhB protein (Uniprot ID: Q9HXI4) was cloned and overexpressed in *E. coli* expression host [BL21(DE3)] using pET28a expression vector with a N-terminal (Histidine)_6_ tag for purification. The recombinant N-(His)_6_-PaSuhB was overexpressed using IPTG induction and was later purified by tandem immobilised metal affinity chromatography (IMAC) (using Ni-Sepharose matrix) and size exclusion chromatography (SEC) (using Superdex-75 matrix). The purity of the PaSuhB protein samples were assessed by its subunit molecular mass (∼30kDa), indicated by the presence of a single band in the SDS-PAGE (12%). Moreover, a sharp single Gaussian SEC elution profile of purified PaSuhB indicates a homogeneously purified protein. (Supplementary Fig. S1). The purified PaSuhB protein showed Mg^2+^ dependent D-*myo*-inositol 1-monophosphate (IPD) phosphatase activity *in vitro* (Supplementary Fig. S2). Importantly, for the crystallization trials, the PaSuhB was purified in presence of 10mM CaCl_2_, to inhibit the hydrolysis of the bound substrate during co-crystallization experiments. For the crystallization of PaSuhB/Ca^2+^/IPD ternary complex, The Ca^2+^ supplemented protein was further concentrated up to 25mg/mL (using membrane filter of 10kDa cutoff) and incubated (for 30 min at ambient temperature) with 5-fold molar excess of substrate (IPD). The apo-form of the PaSuhB protein was crystallized in the absence of the substrate.

The apo and the substrate (IPD) bound forms of PaSuhB protein were crystallized in orthorhombic space group (P2_1_2_1_2_1_) (Table 1). After diffraction data collection, indexing, scaling and merging of the collected data, the molecular replacement using *E. coli* SuhB as template structure (Identity 55.09%) was conducted for initial phase calculation. Iterative cycles of model building and crystallographic refinement was conducted until the final models were in a good agreement with all geometric and stereochemical parameters as well as until a good convergence of crystallographic R factor and R_free_ was attained. After a thorough validation, the coordinates of apo- and substrate (IPD)-bound forms of PaSuhB were submitted to Protein Data Bank (PDB) along with their respective structure factor files. The PDB IDs of PaSuhB apo form is 8WIP (Figure 1A) and PaSuhB/Ca2^+^/IPD bound form is 8WDQ (Figure 1C). The data collection and refinement statistics of these two crystal forms are summarized in Table 1. Both the PaSuhB structures exhibit the dimeric form of the protein and are quite reminiscent with most of the IMPase orthologs ranging from human to bacteria. PaSuhB dimeric structure shows highest similarity to *Bartonella henselae* SuhB (PDB ID: 3LV0) with an overall RMSD value of 0.793Å for all Cα atoms followed by SuhB orthologues from *Escherichia coli* (PDB ID: 6IB8) and *Thermotoga maritima* (PDB ID: 2P3N) with RMSD values (for all Cα atoms) of 1.008 Å and 1.014Å respectively. Conversely the aminos acid sequence similarity of PaSuhB is higher with *E. coli* SuhB (55.09%) followed by *Bartonella henselae* (44%) and *Thermotoga maritima* (40%). Monomeric structures of each dimer (Figure 1B) preserve its classical layered architecture (N-αβαβα-C) as also observed in other proteins of this family including IMPases and FBPases. The two interacting monomers of apo-PaSuhB dimer are structurally very similar to each other (RMSD: 0.126Å). The globular (46X46X35Å) monomeric structure of PaSuhB is guarded by two sets of α helices (α1, α2 and α4, α7) from the top and the bottom. The overall structure is composed of six large alpha helices arranged in separate layers (α1-α2, α5-α6, and α4-α8) with two additional small helix (α3 and α7) and two sets of central beta sheets connected by a long flexible loop. The two sets of central β-sheets, seven in one set (β1 to β7) and five in the second set (β8 to β12) is separated by a layer of alpha helices (α5 and α6). The active site catalytic grove of PaSuhB is situated at the junction point of two central β-sheets and the N-terminus of the small α3 helix. The dipole moment of the α3 helix helps to stabilize the negative charge of the PO_4_^3-^ moiety of the incoming substrate, whereas, the highly conserved metal binding residues reside in the loops connecting α1-α2 (Asp38), β1-β2 (Glu67), β3-α3 (Asp86, Asp89) and the N terminal end of α6 helix (Asp216). The active site of PaSuhB is shared by the dimeric interface of the two interacting monomers and is guarded by a long flexible active site mobile loop (Arg25 to Glu42). Although Ca^2+^ ions were added during the purification of PaSuhB protein, the absence of any large difference density peaks at the active site of apo-PaSuhB suggests the absence of any bound metal ions. Likewise, the metal binding amino acids of the apo-PaSuhB were also not oriented to form a proper metal binding coordination geometry. In the apo-PaSuhB structure, the retraction of the active site mobile loop forming amino acids resulted in the disposition of the open and empty active site of PaSuhB with the solvent accessible area of ∼ 869.35 and 703.87 Å^2^ for chain A and chain B respectively which is comparable to IMPases with stringent substrate specificity (human IMPase and *E coli* IMPases) but is much smaller than the IMPases with promiscuous substrate specificity (dual specific IMPase/NADP(H) phosphatase from Archaea and *S. aureus*). Like, *E. coli* SuhB, in PaSuhB, the increased length and structural kink of α4 helix and its N-terminal loop may be responsible for this structural discrepancy [13]. The dimeric and monomeric organization of IPD bound PaSuhB crystal structure is very similar to the apo form structure (overall Cα RMSD is 0.196Å) except at the active site catalytic cleft and by the differential disposition of active site mobile loop. The average B-factor analysis indicates that the IPD bound crystal structure of PaSuhB (average B-factor 27.64) is more stable than its apo form (average B-factor 40.31). The presence of huge difference density peaks (16.79 and 15.98σ respectively for Ca^2+^ and phosphate of IPD) at the active site of PaSuhB monomers led us to model the bound metal ions (Ca^2+^) and the substrate molecules (IPD). Importantly, the active site of the A chain of PaSuhB/Ca^2+^/IPD complex was found to bind three metal ions whereas, the B chain of the same complex was found to bind two metal ions. However, active site of both these chains is occupied by one molecule of IPD (Figure 1D). Each of these bound metal ions and the substrate molecules could be refined to full (100%) occupancy. The additional presence of the third metal ion at the active site of A chain of PaSuhB/Ca^2+^/IPD complex could be corroborated with the closure of the active site mobile loop resulting in an RMSD of 0.153 Å in between the A and B chains of the PaSuhB/Ca^2+^/IPD complex. Overall, the active site cavity is formed by the amino acid residues including Glu67, Asp86, Leu88, Asp89, Gly90, Thr91, Phe161, Arg162, Arg188, Gly190, Ala191, Ala192, Glu209, Leu212 and Asp216 (Figure 1E and 1F). The first metal ion (Ca^2+^1 in purple colour 2D figure 1E) interacts with the side chains of three highly conserved amino acid residues Asp86, Asp89 and Asp216 at the active site of PaSuhB. The same metal ion also supports the binding of the substrate IPD by interacting with O1 and O8 atoms of the bound substrate. The second metal ion (Ca^2+^2 in purple colour 2D figure 1E) interacts with the side chains of the highly conserved Glu67, Asp86 and the carbonyl oxygen of Leu88 at the active site of PaSuhB. Like the first metal ion, the second metal ion also stabilize the substrate binding by interacting with O8 atom of the IPD. The third metal ion, exclusively obtained in the A chain of the PaSuhB/Ca^2+^/IPD complex, (Ca^2+^3 purple colour in figure 1E) interacts with only two conserved amino acids, the side chains of Asp38 and Glu67. Just like two other active site bound metal ions, the third metal ion also stabilize the binding of the incoming substrate by interacting with O9 atom of the IPD molecule (Figure 1E and 1F). Other than the PaSuhB active site amino acid residues and the bound substrate molecule, each of the metal binding coordination spheres are also comprised of many water molecules. Couple of these water molecules also play pivotal roles in substrate hydrolysis and product release. However, roles of these water molecules in PaSuhB mediated substrate hydrolysis will be described elsewhere. Other than the active site bound metal ions, guanidium groups of Arg162 and Arg 188 stabilize the incoming substrate by formation of hydrogen bonds with O5 and O4 atoms of IPD respectively.

**Table 1:**
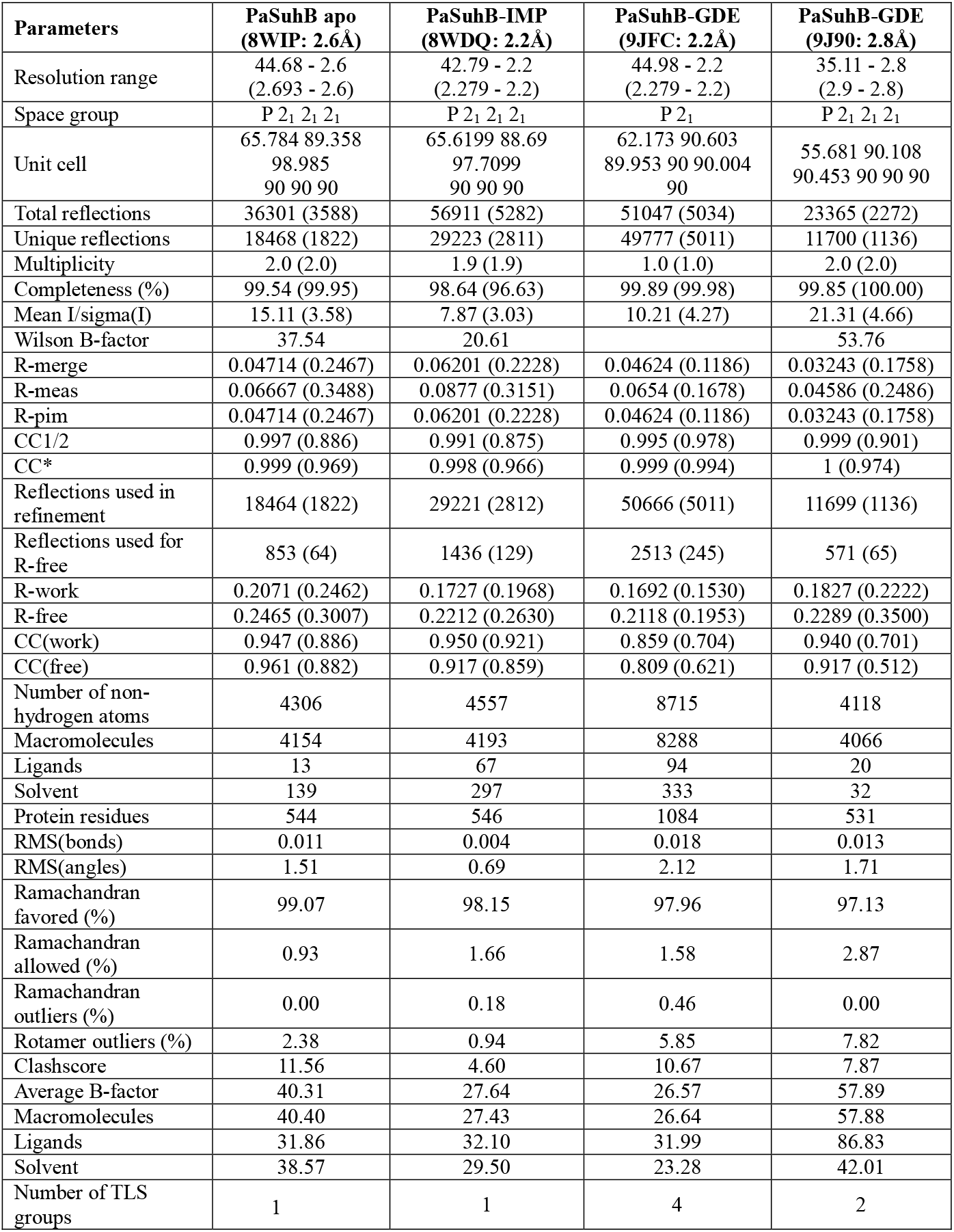
Crystallization parameters for PaSuhB (apo), PaSuhB-IPD and PaSuhB -GDE bound.

| Parameters | PaSuhB apo<br>(8WIP: 2.6Å) | PaSuhB-IMP<br>(8WDQ: 2.2Å) | PaSuhB-GDE<br>(9JFC: 2.2Å) | PaSuhB-GDE<br>(9J90: 2.8Å) |
| --- | --- | --- | --- | --- |
| Resolution range | 44.68 - 2.6<br>(2.693 - 2.6) | 42.79 - 2.2<br>(2.279 - 2.2) | 44.98 - 2.2<br>(2.279 - 2.2) | 35.11 - 2.8<br>(2.9 - 2.8) |
| Space group | P 2<sub>1</sub> 2<sub>1</sub> 2<sub>1</sub> | P 2<sub>1</sub> 2<sub>1</sub> 2<sub>1</sub> | P 2<sub>1</sub> | P 2<sub>1</sub> 2<sub>1</sub> 2<sub>1</sub> |
| Unit cell | 65.784 89.358<br>98.985<br>90 90 90 | 65.6199 88.69<br>97.7099<br>90 90 90 | 62.173 90.603<br>89.953 90 90.004<br>90 | 55.681 90.108<br>90.453 90 90 90 |
| Total reflections | 36301 (3588) | 56911 (5282) | 51047 (5034) | 23365 (2272) |
| Unique reflections | 18468 (1822) | 29223 (2811) | 49777 (5011) | 11700 (1136) |
| Multiplicity | 2.0 (2.0) | 1.9 (1.9) | 1.0 (1.0) | 2.0 (2.0) |
| Completeness (%) | 99.54 (99.95) | 98.64 (96.63) | 99.89 (99.98) | 99.85 (100.00) |
| Mean I/sigma(I) | 15.11 (3.58) | 7.87 (3.03) | 10.21 (4.27) | 21.31 (4.66) |
| Wilson B-factor | 37.54 | 20.61 |  | 53.76 |
| R-merge | 0.04714 (0.2467) | 0.06201 (0.2228) | 0.04624 (0.1186) | 0.03243 (0.1758) |
| R-meas | 0.06667 (0.3488) | 0.0877 (0.3151) | 0.0654 (0.1678) | 0.04586 (0.2486) |
| R-pim | 0.04714 (0.2467) | 0.06201 (0.2228) | 0.04624 (0.1186) | 0.03243 (0.1758) |
| CC1/2 | 0.997 (0.886) | 0.991 (0.875) | 0.995 (0.978) | 0.999 (0.901) |
| CC* | 0.999 (0.969) | 0.998 (0.966) | 0.999 (0.994) | 1 (0.974) |
| Reflections used in refinement | 18464 (1822) | 29221 (2812) | 50666 (5011) | 11699 (1136) |
| Reflections used for R-free | 853 (64) | 1436 (129) | 2513 (245) | 571 (65) |
| R-work | 0.2071 (0.2462) | 0.1727 (0.1968) | 0.1692 (0.1530) | 0.1827 (0.2222) |
| R-free | 0.2465 (0.3007) | 0.2212 (0.2630) | 0.2118 (0.1953) | 0.2289 (0.3500) |
| CC(work) | 0.947 (0.886) | 0.950 (0.921) | 0.859 (0.704) | 0.940 (0.701) |
| CC(free) | 0.961 (0.882) | 0.917 (0.859) | 0.809 (0.621) | 0.917 (0.512) |
| Number of non-hydrogen atoms | 4306 | 4557 | 8715 | 4118 |
| Macromolecules | 4154 | 4193 | 8288 | 4066 |
| Ligands | 13 | 67 | 94 | 20 |
| Solvent | 139 | 297 | 333 | 32 |
| Protein residues | 544 | 546 | 1084 | 531 |
| RMS(bonds) | 0.011 | 0.004 | 0.018 | 0.013 |
| RMS(angles) | 1.51 | 0.69 | 2.12 | 1.71 |
| Ramachandran favored (%) | 99.07 | 98.15 | 97.96 | 97.13 |
| Ramachandran allowed (%) | 0.93 | 1.66 | 1.58 | 2.87 |
| Ramachandran outliers (%) | 0.00 | 0.18 | 0.46 | 0.00 |
| Rotamer outliers (%) | 2.38 | 0.94 | 5.85 | 7.82 |
| Clashscore | 11.56 | 4.60 | 10.67 | 7.87 |
| Average B-factor | 40.31 | 27.64 | 26.57 | 57.89 |
| Macromolecules | 40.40 | 27.43 | 26.64 | 57.88 |
| Ligands | 31.86 | 32.10 | 31.99 | 86.83 |
| Solvent | 38.57 | 29.50 | 23.28 | 42.01 |
| Number of TLS groups | 1 | 1 | 4 | 2 |

**Figure 1:**
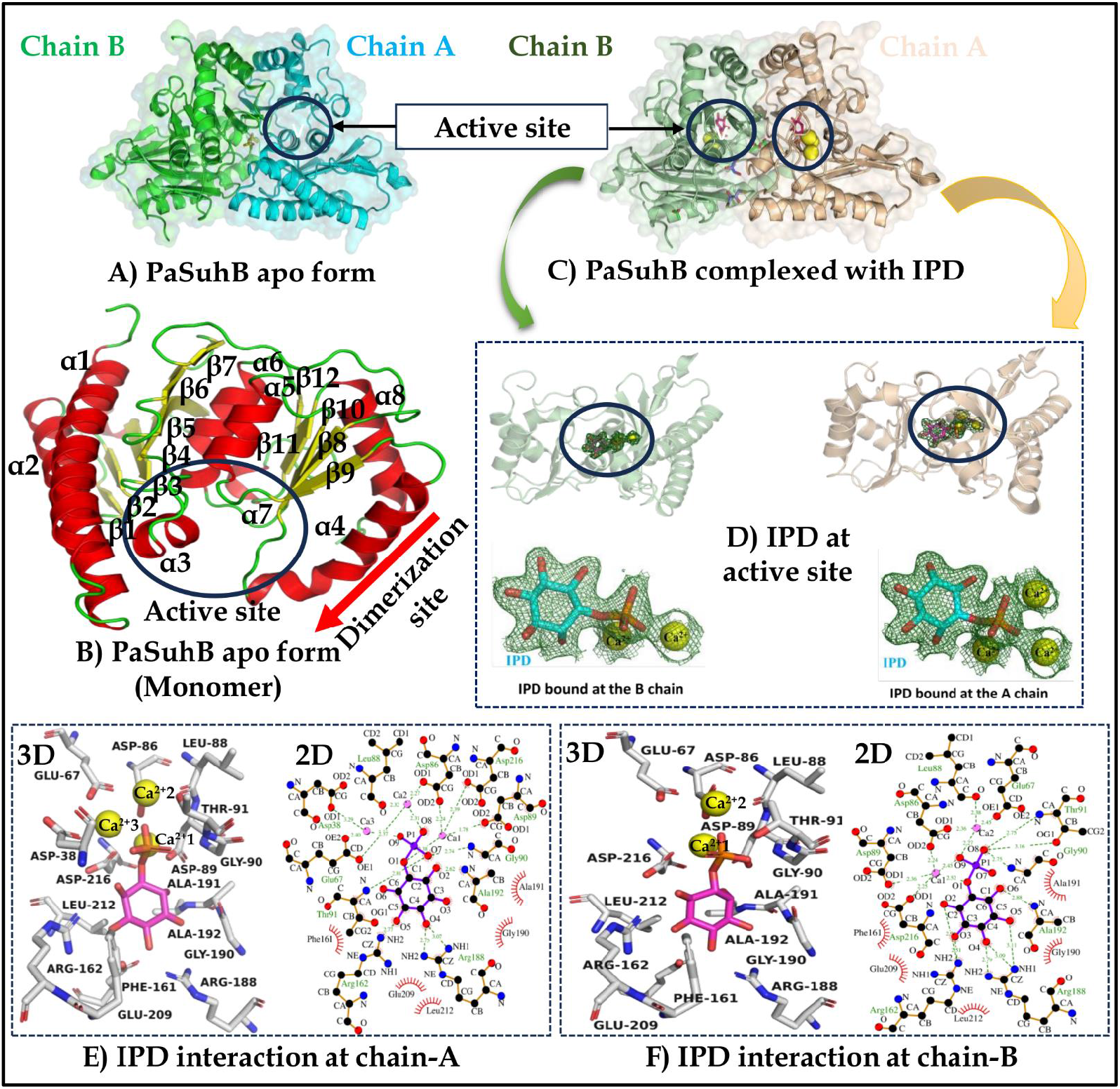
3D structural disposition and substrate interaction profile of PaSuhB apo and IPD bound crystal structures. Figure A) PaSuhB apo structure in dimer form, Figure B) Secondary structural representation of apo PaSuhB monomeric structure, Figure C) PaSuhB complexed with its substrate IPD, two active sites at each monomer shown in circles, IPD shown in magenta while metal ions in yellow colours, Figure D) Represent individual monomers of PaSuhB-IPD complex structure showing 2Fo-Fc map (1σ) of substrate and metal ions, Figure E) Interaction profile of PaSuhB and IPD at chain-A and Figure F) at chain-B active site respectively, residues with red spikes interact with hydrophobic bonds while other with hydrogen bond.

The three calcium ions bound to the A chain of PaSuhB/Ca^2+^/IPD complex have B-factors of 22.72, 34.90 and 37.89 Å^2^ which corroborates their relative stabilities depending on their strength of binding at the PaSuhB active site catalytic cleft. Again, the two calcium ions bound at the B chain of PaSuhB/Ca^2+^/IPD complex have their B-factors of 24.15 and 42.12 Å^2^. Moreover, the substrate IPDs, bound at the chain A and chain B of PaSuhB/Ca^2+^/IPD complex have B-factor of 21.39 and 24.73 Å^2^ respectively. Interestingly, this B-factor analysis of bound metals and substrate at the PaSuhB active site, may suggest that the binding of the 3^rd^ metal ion increases the stability of the 1^st^ and 2^nd^ metal ions as well as the stability of the incoming substrate. Interestingly, unlike the apo-PaSuhB dimer and the B chain of the PaSuhB/Ca^2+^/IPD complex, the active site mobile loop (Arg25 to Glu42) of the A chain adopts a close conformation in order to extend the side chain of Asp38 residue to cater the proper coordination geometry for Ca^2+^3 binding. The closure of the active site mobile loop in chain A of PaSuhB/Ca^2+^/IPD complex resulted in the reduction of solvent accessible surface area of the active site to ∼558.15/329.60Å^2^. The closure of the three metal and substrate loaded PaSuhB active site prevents the exposure of bulk water during the process of catalysis and represents the pre-catalytic state of the protein (Figure 1D cartoon representation and Supplementary Figure 3).

### Pharmacophore based ligand scouting identifies poly-hydroxy carboxylic acids as naturally obtained potential inhibitors of PaSuhB

PaSuhB was previously identified as a pivotal pleotropic virulence regulator of *P. aeruginosa*. Moreover, recent cryo-EM based structural studies explored the involvement of SuhB protein analogues in NusA-dependent ribosome biogenesis/maturation in *E. coli*. In the present work, to identify naturally occurring small molecule-based inhibitor leads of PaSuhB, we leveraged the essential pharmacophoric features of the IPD-bound PaSuhB crystal structure (Figure 2A and 2B). The extracted essential pharmacophoric features (hydrogen bond donor/acceptor, hydrophobic interactions, aromatic interactions, charged interactions etc) of PaSuhB/Ca^2+^/IPD complex were used to virtually screen for naturally existing chemical analogues that preserve these interactions using SwissSimilarity server [35-36]. Intriguingly, polyhydroxy carboxylic acids, especially polyhydroxy aryl/cyclohexyl carboxylates/phosphates or sugar acid lactones were found to bear similar pharmacophoric features of the bound IPD molecule at the active site of PaSuhB dimer (Figure 2C). Initial screening yielded hundreds of molecules; however, we employed structure-guided rational screening and have shortlisted only the top fifteen poly-hydroxy carboxylates for further *in-vitro* validation experiments (Supplementary Table S1). Polyhydroxy phosphates were excluded as they may be potential alternative substrates of PaSuhB or any other non-specific cellular phosphatases.

**Figure 2:**
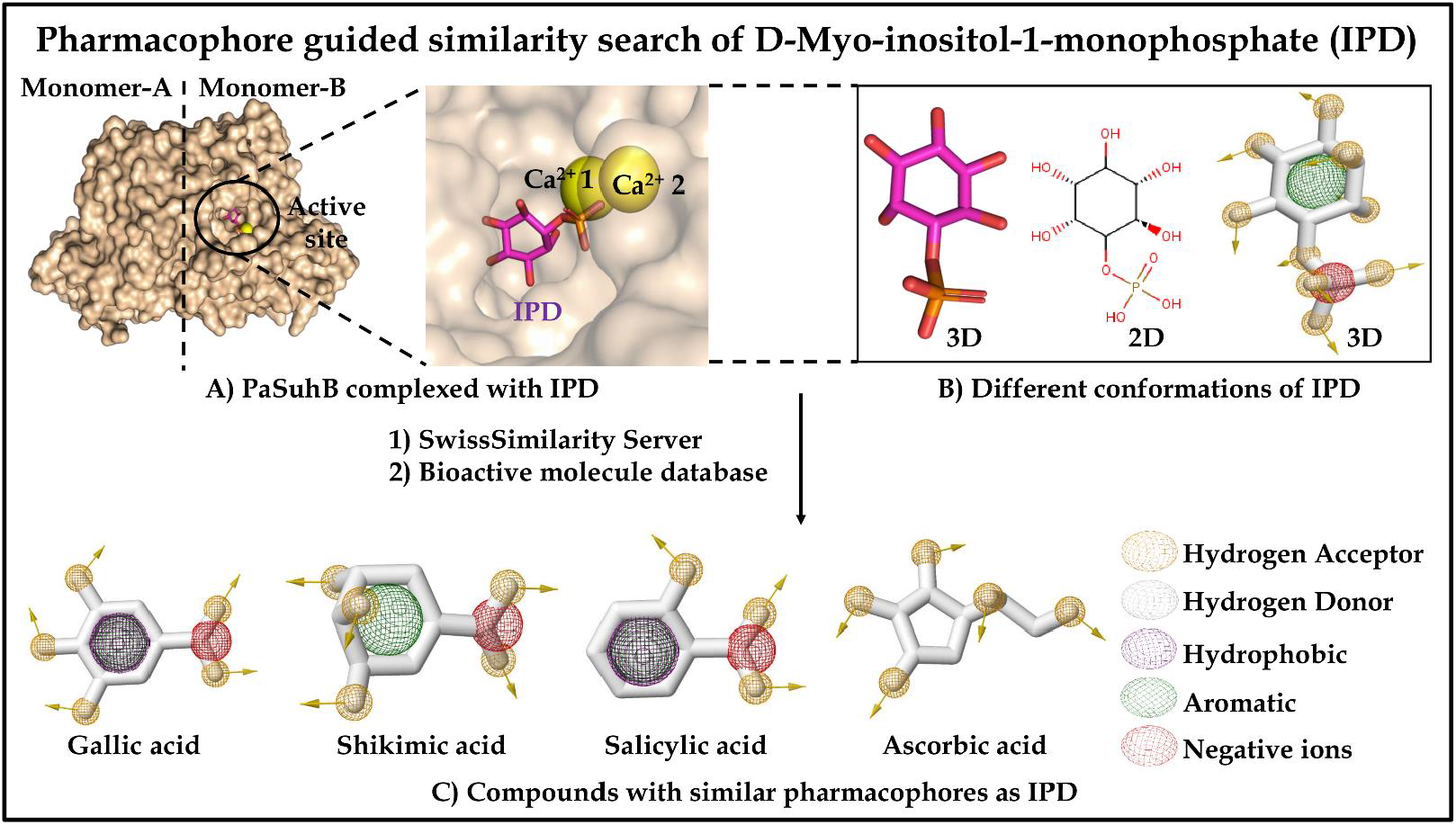
Pharmacophore guided similarity search of IPD. Figure A) Surface view of PaSuhB dimeric structure in complex with IPD (magenta) and metal ions (yellow) at active site, Figure B) Different representation of IPD structure to visualize 3D and 2D conformations along with its pharmacophoric details, Figure C) Some of the similar compounds representing similar pharmacophores as IPD.

### Gallic acid binds PaSuhB protein at low micromolar concentrations and inhibits the enzymatic activity of PaSuhB as a non-competitive inhibitor

Pharmacophore guided inhibitors screening provides an array of small molecule based natural compounds with predicted potential to interact at PaSuhB active sites, which is then followed by validation of these compounds through investigation of PaSuhB enzyme activity inhibition, utilising colourimetry-based malachite green phosphate assay kit [MAK307 Sigma-Aldrich]. Enzyme activity inhibition study reveals that one of the naturally existing molecules, GDE exerts the most potent inhibition among those fifteen shortlisted polyhydroxy/polyphenol carboxylate molecules (Supplementary Table S1). GDE shows concentration-dependent inhibition of PaSuhB enzyme activity with IC_50_ value of approximately 1.34mM (Figure 3A). Enzyme kinetics-based double reciprocal plot study was conducted to explore the mode of GDE mediated PaSuhB IMPase activity inhibition, which reveals that GDE shows a noncompetitive mode of inhibition as extrapolated graph in presence of with and without inhibitors intersecting at X axis. Furthermore, these results were supported by values of kinetic parameters like K_m_ and V_max_, as in case of typical noncompetitive inhibition, the substrate’s K_m_ remains constant and V_max_ decreases upon increasing inhibitor concentration. Similar observations were reported in case of GDE based PaSuhB inhibition (Figure 3B). Again, the noncompetitive mode of enzyme inhibition indicates that the formation of Enzyme-Substrate-Inhibitor (ESI) complex as well as the Enzyme-Inhibitor (EI) complex is equally possible.

**Figure 3:**
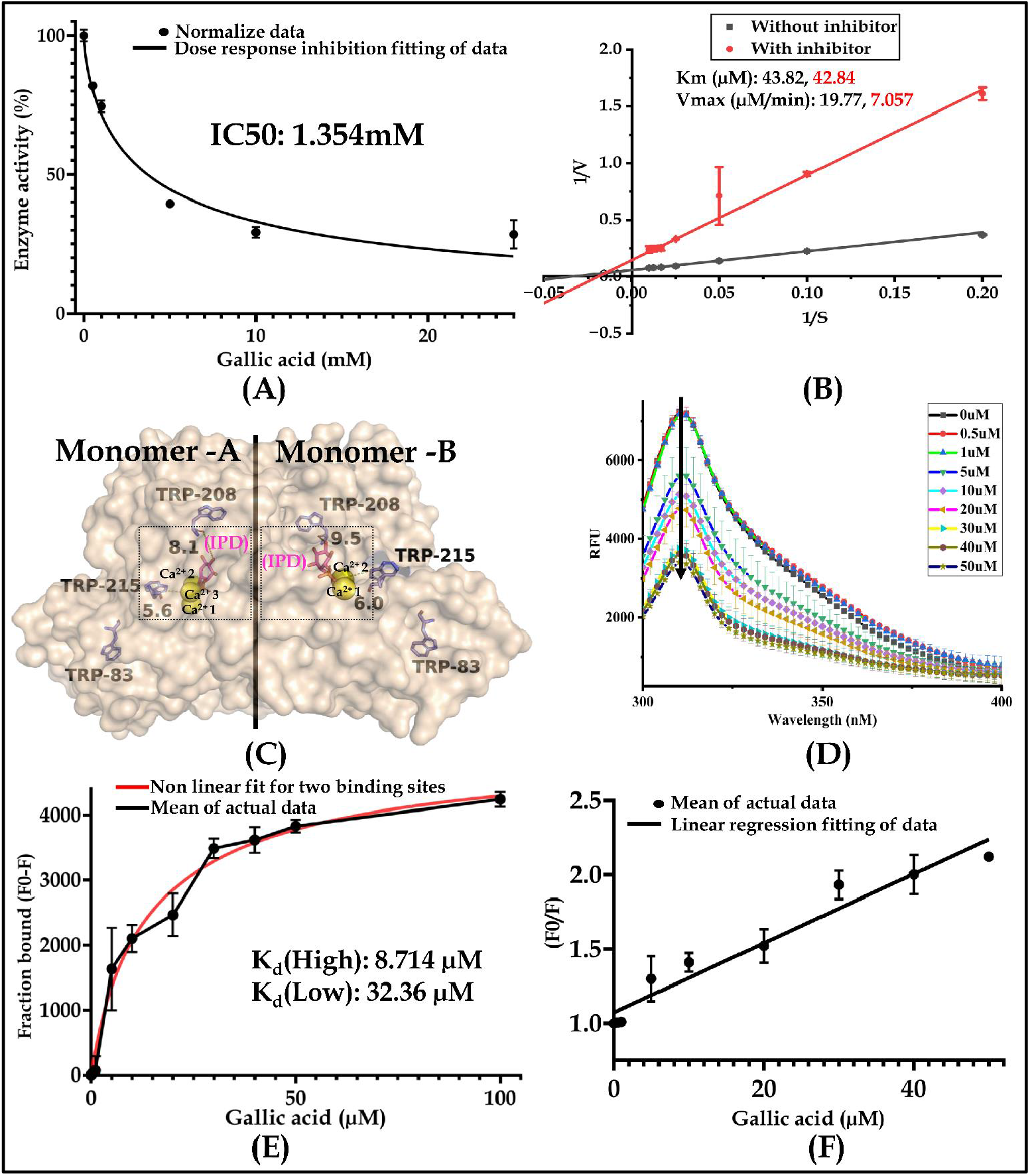
GDE based PaSuhB enzyme activity inhibition and fluorescence-based protein-ligand interaction study. **Figure** A) GDE concentration dependent inhibition of PaSuhB IMPase activity, Figure B) Double reciprocal plot analysis of PaSuhB enzyme kinetic data in presence and absence of GDE. Figure C) Surface view of IPD bound PaSuhB structure to show zoomed view of tryptophan in near vicinity of active sites (shown in dotted rectangle), IPD (magenta), metal ions (yellow) and tryptophan (blue). Figure D) GDE concentration dependent tryptophan fluorescence quenching study, Figure E) GDE-PaSuhB binding isotherm analysis to represent number of binding sites and their respective binding affinities, Figure F) Stern-Volmer plot of PaSuhB-GDE interaction.

This suggests the possibility of more than one mode of GDE binding at the active site of PaSuhB. Furthermore, to evaluate the number and mode of GDE binding at PaSuhB active site, tryptophan fluorescence quenching-based titration of GDE binding to PaSuhB was conducted. Importantly, each PaSuhB monomer contains three tryptophan residues (Trp83, Trp208 and Trp215) in the close vicinity of active sites (Figure 3C). GDE exerts concentration-dependent quenching of PaSuhB intrinsic fluorescence (Figure 3D), which confers the interaction between PaSuhB with GDE. Accordingly, the analysis of the Langmuir isotherm reveals the presence of two plausible GDE binding sites with affinities in the low micromolar ranges 8.714 and 32.36µM respectively (Figure 3E). Furthermore, partial downward deviation of the linear fit of Stern-Volmer plot further indicates the presence of multiple GDE binding sites and a plausible negative cooperativity amongst these binding sites (Figure 3F). Overall, the *in-vitro* biochemical and biophysical data of GDE interaction with PaSuhB reveal multimodal GDE binding at PaSuhB active site at low micromolar ranges while the GDE mediated inhibition of PaSuhB enzymatic activity occurs at low millimolar ranges.

### Crystal structure of PaSuhB/GDE complex unravels the two distinct modes of GDE binding, imperative of the GDE mediated noncompetitive mode of inhibition of PaSuhB enzymatic activity

The biochemical and biophysical studies of PaSuhB-GDE interaction indicate two distinct modes of GDE interactions with PaSuhB. To visualize the exact mode of binding of GDE with PaSuhB as well as to excavate the molecular mechanism of GDE inhibition, we solved the crystal structures of PaSuhB-GDE binary complexes. PaSuhB/GDE complex was prepared by gently mixing 10-fold molar excess of GDE with the purified Mg^2+^ supplemented homogeneously purified PaSuhB protein and incubating the mixture at room temperature for 30 minutes before crystallisation drop setting. The PaSuhB-GDE complex was crystallised in similar crystallisation conditions of apo-PaSuhB and the PaSuhB-Ca^2+^-IPD complex. However, unlike apo-PaSuhB and PaSuhB-Ca^2+^-IPD complex, PaSuhB-GDE complex reveals two distinct crystal packings. The monoclinic (P2_1_) form of PaSuhB-GDE complex diffracted to higher resolutions (2.2 Å); while the orthorhombic (P2_1_2_1_2_1_) form diffracted up to 2.8Å. Both of these two crystal structures were phased using Molecular Replacement by using apo-PaSuhB as the template; their finally refined atomic coordinates along with structure factor files were submitted to PDB (PDB IDs are 9JFC and 9J90 for P21 and P212121 space group bearing structures respectively). The crystallographic data collection and refinement parameters of these two structures are also summarised in Table1. Interestingly, both of these two different forms of PaSuhB-GDE complexes demonstrate the presence of active site-bound GDE. However, neither of these two structures shows difference density peaks which correspond to the bound metal ions at the PaSuhB active site. Overall, both of these two structures show a high degree of structural resemblance with each other corroborated by comparative RMSD analysis. The RMSD value of low-resolution structure’s chain A with high resolution’s chain A, E, I and M, are 0.253, 0.287, 0.266 and 0.288Å respectively while low-resolution structure’s chain B with high resolution’s chain A, E, I and M, are 0.260, 0.245, 0.231 and 0.255Å respectively. Similar observations are obtained with RMSD analysis at dimeric level as the RMSD value of low-resolution structure AB chain to high resolution AE and IM chains are 0.488 and 0.399Å respectively. Both of these two crystal structures are very similar to PaSuhB apo forms of crystal structure as RMSD values of apo-PaSuhB AB dimers are 0.477Å with low-resolution AB dimer while 0.246 and 0.279Å are with high resolution’s AE and IM dimers respectively. Unlike the high-resolution crystal structure of PaSuhB/GDE complex (solved in P2_1_ space group) which shows the presence of GDE in the active site of each protomer, the lower resolution structure (solved in P2_1_2_1_2_1_ space group) of the PaSuhB/GDE complex demonstrates the presence of GDE in the active site of only one chain of the PaSuhB dimer. (Protein-ligand interaction of the lower-resolution structure is given in Supplementary Figure S4). More interesting findings were observed in the high-resolution crystal structure of PaSuhB-GDE complex intriguingly, this structure was solved in an unusual asymmetric tetrameric form (Figure 4A). Although classical Fructose 1,6 bisphosphate phosphatase mimicking tetrameric structures have been previously encountered in IMPase orthologs present in hyperthermophiles eubacterial (PDB ID: 2P3N) and archaeal (PDB ID: 2PCR) species. In most of these high temperature dwelling archaea and eubacterial IMPase orthologs also show promiscuous FBPase activity [39]. However, the structure of GDE bound PaSuhB tetramer is novel in terms of the interaction in between two PaSuhB dimers.

**Figure 4:**
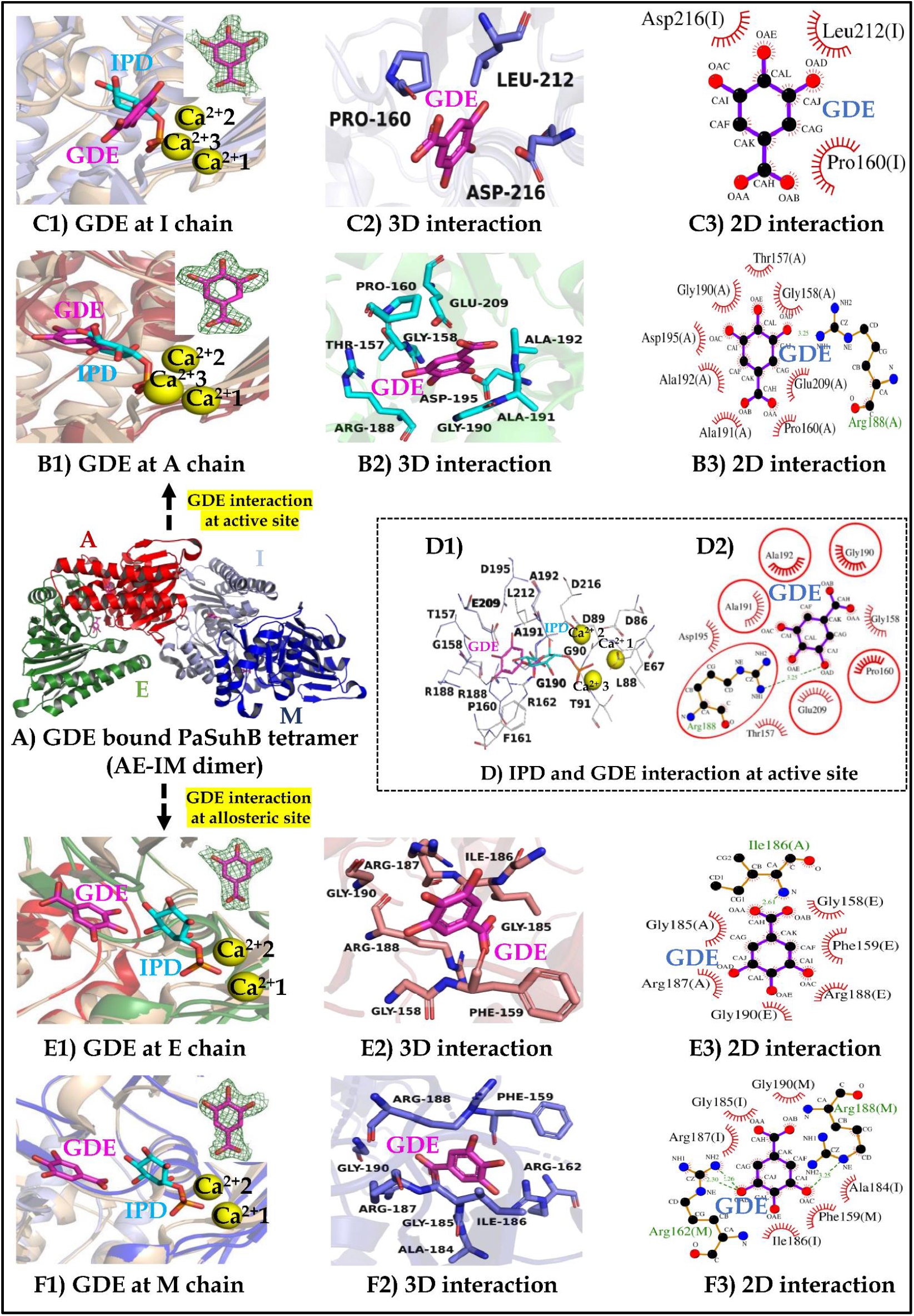
GDE bound PaSuhB crystal structure and their interaction profile. Figure A) PaSuhB complexed with GDE, crystal structure in tetrameric form. Tetramer is composed of two homodimers (AE and IM dimer), in fist dimer A chain (red) and E chain (Green) while in second dimer I chain (light blue) and M chain (Dark blue), ‘A’ monomer of AE dimer and ‘I’ mopnomer of IM dimer inetract to form tetramer. Figures B1, C1, E1 and F1 represents active site of GDE bound PaSuhB overlaped with IPD bound PaSuhB to show relative position of GDE with respect of IPD. Figure B1) and C1) represent GDE interaction at chain A and chain I active sites while figure E1) and F1) represent GDE interaction at dimerization (allosteric sites), Electron density map (2Fo-Fc) of GDE in figure B1, C1, E1 and F1 shown in green colour (countoured at 1σ), GDE (magenta) and IPD (cyan). B2 and B3 represent 3D and 2D interaction profile of chain A while C2 and C3 represents chain I GDE. Figure D) Investigation of identical residues involved in IPD and GDE interaction at PaSuhB active site, D1) 3D representation D2) 2D interaction, encircled residies presents identical residues. Figures E2 and E3 represents 3D and 2D interaction of GDE at AE dimerization surface while F2 and F3 at IM dimerization surface. In figures B3, C3, D2, E3 and F3 residues with red spikes represents hydrophobic interaction while others form hyderogen bonding.

The tetramer of PaSuhB-GDE complex (PDB ID: 9JFC) is formed by two homodimers, one composed of A and E chains and another one with I and M chains. The tetramer was asymmetrically formed via protein-protein interaction in between A chain of AE dimer and I chain of IM dimer. Secondary structural organization of both these two interacting dimers are overall similar to the dimer of PaSuhB IPD-bound (PDB ID: 8WDQ) and apo-PaSuhB (8WIP) structures with few localised structural differences. While superimposed on the dimers of PaSuhB-IPD complex, the tetramer forming AE and IM dimers of the PaSuhB-GDE complex have RMSD (all Cα atoms) of 0.278Å and 0.282Å, respectively. Once again, these two interacting dimers of the tetrameric assembly also demonstrate interesting structural differences resulting in the Cα atoms based RMSD of 0.296Å. The main structural differences between the two interacting dimer stems arises from the structural disposition of α4 helix and its preceding loop on each PaSuhB monomer, which indicates a different pattern of GDE binding (Supplementary Figure 5A1-A3).The tetrameric structure of the PaSuhB-GDE complex (average B-factor 26.57 Å^2^) demonstrates the presence of four active site bound GDE molecules (one at each monomer), however, no metal ions could be detected at the active sites of any monomer. Each dimer of tetrameric structure contains two molecules of bound GDE; interestingly, these two bound GDE molecules have two distinct binding modes at their corresponding monomeric binding sites, which also supports the finding of two distinct modes of GDE binding through fluorescence quenching based titration data (Figure 3E) as well as the noncompetitive mode of GDE inhibition of PaSuhB IMPase activity (Figure 3B). Importantly, the interacting monomers of each dimeric assembly of the tetrameric PaSuhB-GDE complex harbour the bound GDE molecules in four different binding environments. While superimposed with IPD bound PaSuhB dimeric structure, the bound GDE molecules were either found to occupy the IPD binding sites (Fig. 4B, Fig. 4C) [as was found in A chain and I chain and thereby exert the formation of enzyme-inhibitor (EI) complex, which may hinder the binding of incoming substrate] or was found to occupy a novel α4 helix proximal binding site close to the PaSuhB dimeric-interface, away from the IPD binding site (Fig. 4E, 4F) [as was found in E chain and M chain and thereby exert the formation of enzyme-substrate-inhibitor (ESI) complex, which may not hinder the binding of incoming substrate but may hinder the process of catalysis or product release]. All the bound GDE molecules were refined with their full occupancy at the respective binding sites, and the difference density-based omit maps, contoured at 1σ clearly indicates their unambiguous presence at each binding sites of PaSuhB-GDE complex (Fig.4). To our knowledge, for SuhB proteins (and other IMPase orthologs), the discovery of a small molecule-based ligand binding site at the α4 helix proximal PaSuhB dimeric interface is novel. The active site GDE binding residues of PaSuhB are Thr157, Gly158, Pro160, Arg188, Gly190, Ala191, Ala192, Asp195 and Glu209 at Chain A bound GDE (Figure 4B2 and B3) and Pro160, leu212 and Asp216 at chain I bound GDE (Figure 4C2 and C3). Among these nine amino acids (Figure 4B3) only Arg188 form hydrogen bond with hydroxyl group of C3 ring carbon of bound GDE while rest of the residues involved in hydrophobic interactions. Overlapping amino acid residues of PaSuhB which interacts both with GDE and IPD are given in Figure 4D which indicates that GDE can occupy the identical binding site as the substrate IPD. Overall, six interacting amino acid residues, Pro160, Arg188, Gly190, Ala191, Ala192 and Glu209 are common between GDE and IPD interactions (Figure 4D2). Binding of GDE at the PaSuhB active site and engagement of common amino acid residues between IPD and GDE binding corroborates that GDE competitively inhibit PaSuhB enzyme activity.

The PaSuhB-GDE interaction profile at the novel ligand binding site demonstrates GDE binding at the interface of A and E protomer chains (Figure 4E1). At this site, the bound GDE interacts with Gly185, Ileu186, and Arg187 of chain-A while Gly158, Phe159, Arg188 and Gly190 are from chain-E. (Figure 4E2 and E3). Similarly, at the I and M protomer chain interface (Figure 4F1), GDE interacts with Ala184, Gly185, Ileu186 and Arg187 of I chain while Phe159, Arg162, Arg188 and Gly190 are from M chain (Figure 4F2 and F3). At this novel ligand binding site, the PaSuhB-GDE interaction delineates the most insightful observation, which is the interaction of bound GDE with Arg188 residue. Importantly since the identical Arginine residue in *E. coli* SuhB regulates the monomer-dimer equilibrium, which is crucial to the regulatory activity of the SuhB protein in *E. coli in vitro* and *in vivo* [24]. Importantly, the amino acid residues that are involved in the stabilisation of the dimeric interface of PaSuhB were also found to be involved in the formation of the novel binding site of GDE (Supplementary Figure S6). This observation may open up the possibility of avenues where small-molecules may regulate the oligomerisation of PaSuhB monomers to regulate its extragenic suppressor activity *in vitro* and *in vivo*. Collectively, the detailed interaction analysis of the bound GDE molecules found in the tetrameric PaSuhB-GDE complex crystal structure strongly validates our *in vitro* biophysical and biochemical experimental findings, such as the presence of more than one binding site/mode of GDE and the noncompetitive mode of PaSuhB IMPase activity inhibition by GDE.

### GDE binding at the novel ligand binding site of PaSuhB leads shortening of α4 helix to promote higher order oligomerization of PaSuhB and disrupts its pleotropic regulatory functions

In mesophilic eubacteria, the highly conserved SuhB orthologs play pleotropic regulatory functions, which include cold sensitivity, exopolysaccharide biosynthesis, biofilm formation, and antibiotic resistance [18-25]. Recently *E.coli* SuhB was also discovered to play crucial role in ribosomal RNA syntheis, folding and maturation as being part of rRNA transcription antitermination complex via interacting with NusA, NusG and RNA polymerase (Supplementary Figure S7). The high-resolution crystal structure of *E.coli* SuhB dimer with the AR2 domain of NusA protein as well as the cryo-EM based structure of *E.coli* transcription anti-termination complex indicates that the dimeric form of the SuhB protein is involved in these interactions, which facilitates the binding of NusG/E dimer and NusA at processive rRNA sites. An earlier study suggests that the monomeric form of *E.coli* SuhB protein is necessary to interact with RNA polymerase to perform its regulatory role as extragenic supressor [22-24]. Once again, the dimeric form of these family of proteins is necessary to exert the phosphatase activity as their active site is shared by the two interacting monomers [42]. All these studies indicate that to be an active enzyme or active regulators of transcription and translation, the SuhB orthologues should exist in in monomeric and/or dimeric forms and these two forms of the enzyme should be in dynamic equilibrium as per the need of the host bacterial cell. Importantly, the sequence and the structural alignment with *E.coli* SuhB suggests, PaSuhB is highly similar to its *E.coli* counterpart (Supplementary Figure S8). Moreover, structural alignment of apo-PaSuhB dimer with the *E.coli* SuhB dimer-NusA (AR2 domain) complex suggests the high degree of structural preservation of NusA binding site in PaSuhB dimer (Supplementary Figure S9A). Likewise, the SuhB binding sequence in *E.coli* and *P.aeruginosa* NusA AR2 domain is also highly conserved (Supplementary Figure S9B). Therefore, as it is observed in the cryo-EM structure of *E.coli* transcription anti-termination complex, an analogous role of PaSuhB might also be plausible in the context of the earlier reported pleiotropic regulatory functions of PaSuhB. The alteration of oligomeric status of the PaSuhB, especially promoting higher order oligomerization from dimeric structure could be a promising strategy to abolish its regulatory functions. Interestingly, the crystal structure of PaSuhB-GDE complex indicates that binding of GDE at the novel ligand binding sites (at dimeric interface) cause drastic structural deformity at α4 helix and its preceding loop. The detailed comparative structural analysis reveals that the length of α4 helix is reduced from 28-29Å to 21-22Å (Figure 5A for AE dimer and 5B for IM dimer for high resolution and supplementary figure S5 B1, B2 and B3 for low resolution structures) as indicated by the structural alignement of apo-PaSuhB and PaSuhB-GDE bound structures. The critical assessment of 3D strcutures around α4 helix indicates that to compensate the reduced size of α4 helix the sizes of preceding loop and succeding β sheet are increased. Since the substrate IPD only interacts at the active site catalytic cleft while the inhibitor GDE interacts at active site as well as at the dimerization site proximal novel ligand binding site,hence we corroborate that the structural deformity in the GDE bound PaSuhB resulted from the GDE intercation at both sites. Two dimer of GDE bound high resolution PaSuhB crystal structure shows different levels of structural deformity as compared to the apo-PaSuhB structures, which is due to differneces in the GDE binding strengths; in AE dimer one GDE binds at the active site and other near the dimeric interface while in the IM dimer, the active site interaction is very poor, only GDE interaction at the dimeric interface is prevalent (Supplementary Figure S5).

**Figure 5:**
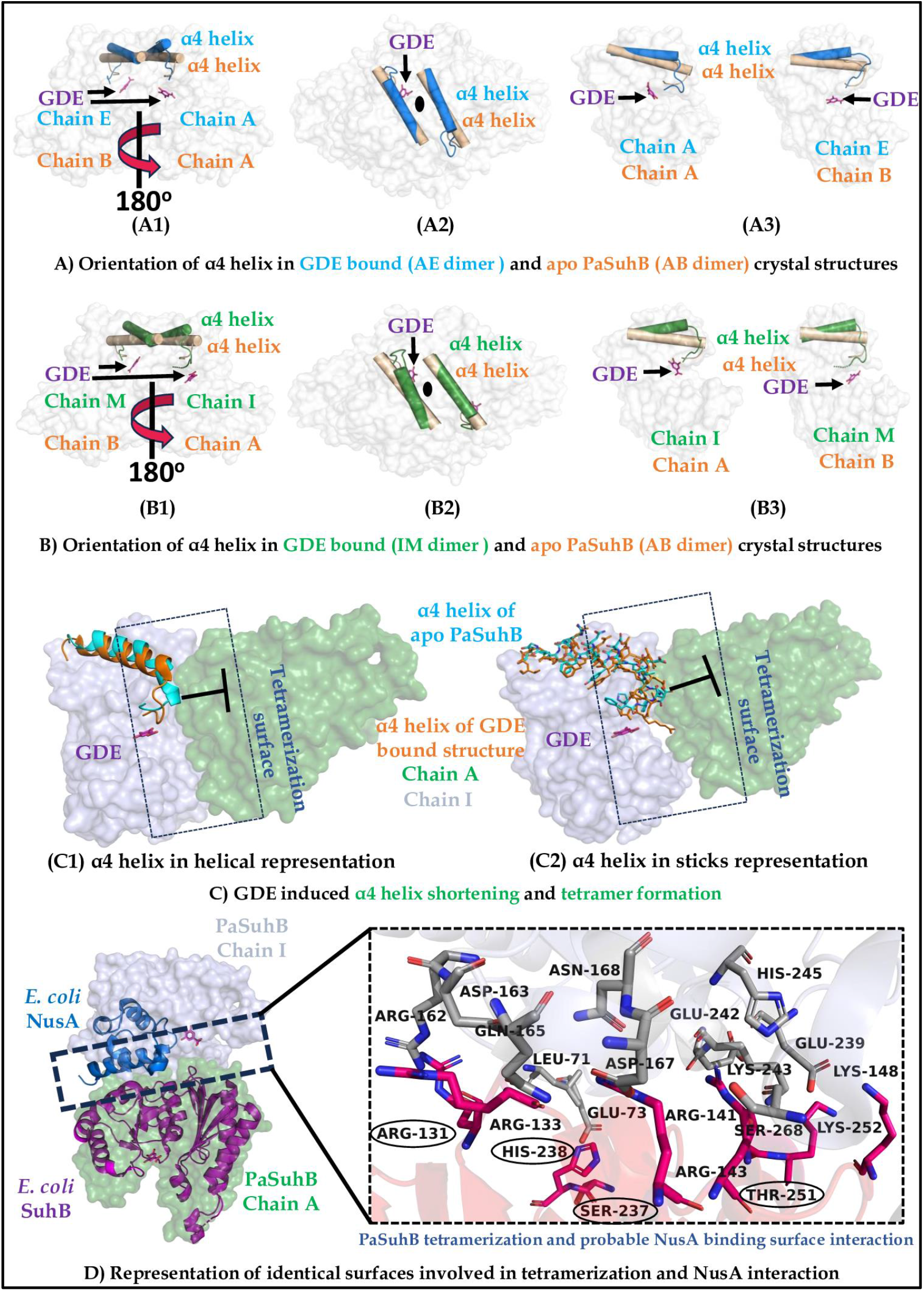
Impact of GDE binding at PaSuhB dimerization site on σ4 helix length, twist angle and oligomerization status of PaSuhB. Figure A) and Figure B) Comparative representation of σ4 helix length and twist angle in AE (cyan) and IM (green) dimers of PaSuhB GDE complexes respectively with respect to PaSuhB apo structure (orange) separately in each dimer, A1 and B1 (top view), A2 and B2 (side view) while A3 and B3 (individual monomers). Figure C) Representation of tetramerization surfaces and clash of α4 helix, PaSuhB GDE bound chain A (green) and chain I (light blue), PaSuhB apo (orange helix) and PaSuhB GDE bound (cyan helix), clashes position is shown with clash sign. Figure D) Representation of identical surfaces and interaction residues between NusA interaction and tetramerization, left panel show the identical surface (shown with dotted rectangle) involved in *E. coli* suhB (magenta cartoon)-NusA (sky blue cartoon) and PaSuhB tetramerization chain I (light blue surface)-chain A(green surface), while right panel shows residues involved in tetramerization, black in circled residues are identical in tetramerization and NusA interaction corroborates common binding site.

From the figures 5A1 and 5B1 it is clearly visible that not only the size of α4 helix but also their relative twist angles at the dimerization interface is altered due to GDE interaction. Thus in the PaSuhB-GDE complex, the IM dimer is more structurally distant and comparatively unstable as compared to the AE dimer as indicated by the overall B-factor analysis (Supplementary Figure S10 A1, B1 and C1). We have also evaluated overall charge distribution, and the results indicate that due to GDE interaction, dimerization interface exposed with slightly higher positive charge than apo-PaSuhB structure (Supplementary Figure S10 A2, B2 and C2). Most importantly, the GDE mediated drastic secondary structural alteration (α4 helix shortening and change in its twist angles) of PaSuhB at its novel ligand binding site also found to alter the acessibility of plausible NusA interaction site of PaSuhB. Hence, we examine all the amino acid residues, their associated B-factors and the charge distribution patterns which might be involved in NusA-AR2 domain interaction. Our observations suggest that few residues like Glu24, Arg112, Arg140, His238 and Lys255 show high degree of conformational alteration in GDE bound structures (Supplemetary Figure S11A). However, residual B-factor and charge distribution analysis of the plausible NusA-AR2 domain binding surfaces of PaSuhB suggest GDE binding did not alter charge distribution of the putative NusA-AR2 domain binding surface of PaSuhB, in fact, B-factor analysis reveals that the putative NusA-AR2 binding region appears relatively stable in GDE bound structure of PaSuhB. However, the reduced B-factor of putative NusA binding amino acids in GDE bound PaSuhB may result from the overall higher resolution of the PaSuhB-GDE complex (2.2Å) compared to the apo-PaSuhB (2.6Å) (Supplemetary Figure S11 B and C). Intriguingly, the analysis of the tetrameric surface of PaSuhB-GDE complex suggests the crucial role of α4 helix and its preceding loop to regulate the tetramer formation in PaSuhB. In apo-PaSuhB dimer the extended length (by 6-7Å) and structural kink of α4-helix impedes the formation of tetramer by imparting steric clashes with the tetramer-forming residues of the neighbouring dimer (Figure 5 C1 and C2). Conversely, PaSuhB-GDE bound structure has shorter and upward shifted α4 helix, which facilitates the spatial accommodation of the tetramer stabilising residues of the neighbouring dimer and promotes higher order oligomer formation. These findings underscore the pivotal role of the α4 helix and its preceding loop in mediating PaSuhB self-oligomerisation, representing a previously unreported novel mechanism of SuhB (and IMPase) oligomerisation which may regulate its biological functions as extragenic suppressor. Moreover, we have analysed the shortened α4 (from AE dimer) interacting tetramerisation surface of the neighboring dimer (IM dimer) of PaSuhB-GDE complex (Figure 5D). Interestingly the putative NusA-AR2 domain binding region of the neighboring dimer (IM dimer) is involved in tetramer formation by interacting with shortened α4 helix and its preceding loop from the AE dimer (Figure 5D). The residues that are common in stabilization of PaSuhB tetramerisation surface and the putative NusA-AR2 domain binding surface are Arg131, His238, Lys252 and Lys255 (indicated by black circles in Figure 5D). The stability of PaSuhB tetramer is also predicted by PDBePISA server, which suggests that the area of tetramerisation interface between chain A of AE dimer and chain I of IM dimer is approximately 991.2 Å^2^ which indicates considerably stable interaction. The overall structural stability of PaSuhB functional dimers of the tetrameric PaSuhB-GDE complex were observed with molecular dynamic (MD) simulation study, where simulation descriptors [like overall Root Mean Square Deviation (RMSD) of Cα atoms, Root Mean Square Fluctuation (RMSF) of amino acid residues, overall Radius of gyration (Rg) and Solvent Accesible Surface Area (SASA) analysis] are compared with the apo-PaSuhB dimer. The descriptors, such as RMSD (Figure 6A) and Rg (Figure 6C), except SASA (Figure 6D), reveal higher quantitative values, which means the structure is less compact compared to the PaSuhB apo structure. Moreover, RMSF analysis (Figure 6B) indicates higher fluctuation in the GDE interacting residues at the active site, while rest of the residues getting stabilized. These MD simulation and crystal structure based observations suggest that the conformational flexibility and shortening of α4 helix imparted by GDE binding at novel ligand binding site of PaSuhB tends to stabilize the overall structure by asymmetrically interacting with self-dimers, resulting in the tetramer formation. However, these observations could be resulted from crystallisation artefact; therefore, to further validate these findings, we performed the Dynamic Light Scattering (DLS) analysis to measure the hydrodynamic diameter of PaSuhB in presence of GDE. Interestingly, the DLS results (Figure 6E and 6F) reveal that in the absence of GDE PaSuhB shows a hydrodynamic diameter ∼5 nm while in the presence of GDE it is doubled to become approximately ∼10nM. The doubling of the hydrodynamic radius of PaSuhB in the presence of GDE fortifies the terameric structure of PaSuhB-GDE complex obtained by X-ray diffraction crystallography. Therefore, in PaSuhB, the involvement of common amino acid residues participating in NusA-AR2 domain interaction as well as the tetramer formation in the presence of GDE, suggests GDE may inhibit the pleotropic extragenic suppressor role of PaSuhB by plausibly disrupting its interaction with NusA and other components of transcription anti-termination complex as observed in *E.coli*.

**Figure 6:**
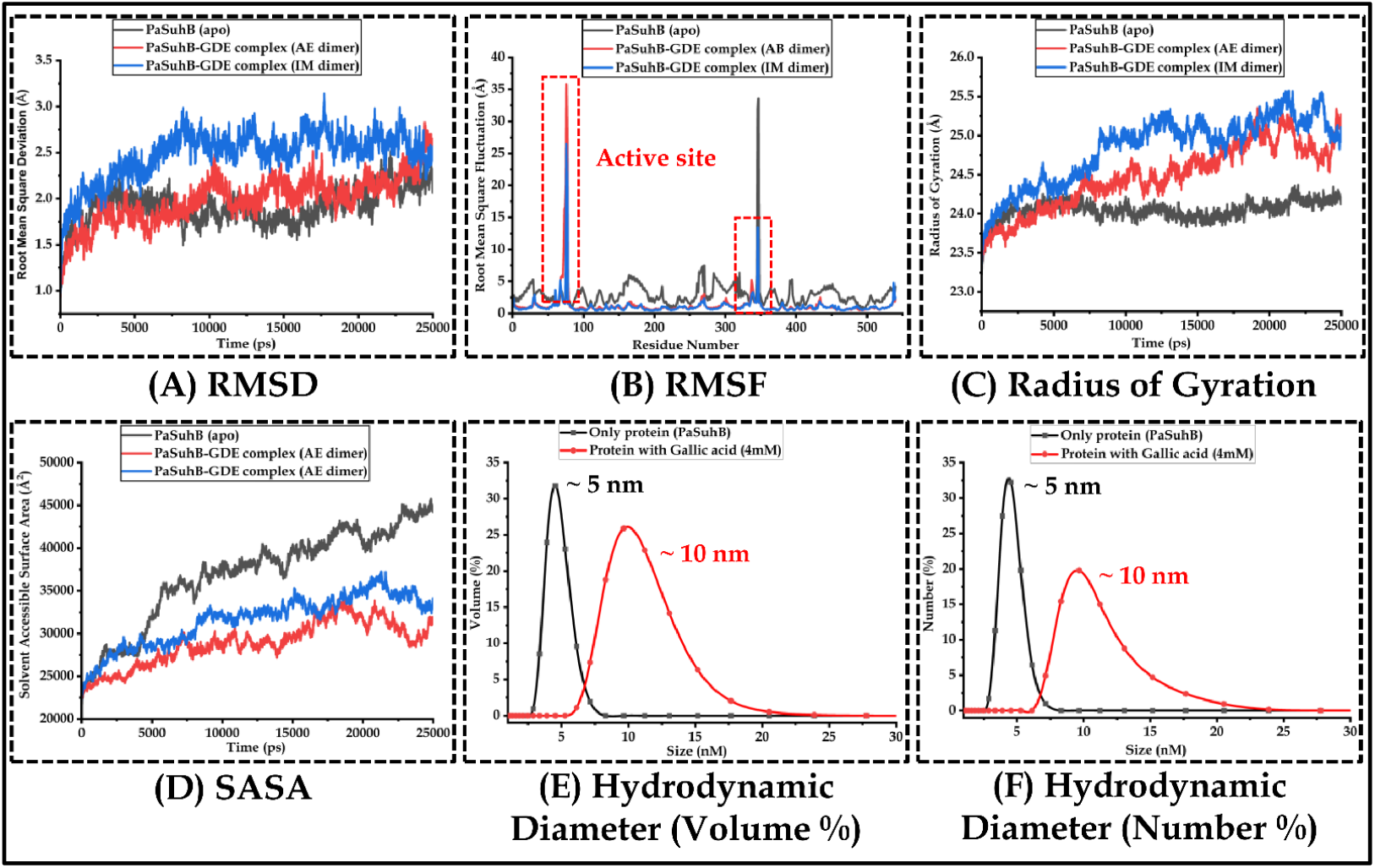
*In-silico* molecular dynamic simulation and in solution DLS study of PaSuhB-GDE interaction. Evaluation of simulation trajectories of PaSuhB apo structure and GDE complex A) Cα RMSD, B) RMSF, C) Radius of gyration, and D) SASA. Measurement of hydrodynamic diameter of PaSuhB in presence of GDE with respect to E) volume% and F) Number%.

### GDE at low concentrations inhibits the swimming motility of *P.aeruginosa* and induces cold sensitive growth phenotype in Gram negative bacteria

Earlier studies demonstrated that PaSuhB deletion mutant exerts an avirulent phenotype in mouse acute infection model, loss of cytotoxicity to the lung cells, decreased expression of T3SS, inhibition of swimming motility and pyocyanin production [25-27]. Furthermore, PaSuhB *E.coli* ortholog null mutants show cold cold-sensitive growth phenotype however, the cold sensitive growth of *E. coli* was not related to SuhB’s IMPase activity [24].

In the present study, the crystallographic as well as biochemical and biophysical data strongly suggests, GDE may inhibit the enzymatic activity as well as extragenic suppressor activity of PaSuhB. These results prompted us to test the effect of GDE specifically on *P.aeruginosa* as well as on an analogous SuhB bearing bacteria, *E.coli*. Firstly, the effect of GDE on the growth of *P.aeruginosa* was estimated; GDE shows concentration-dependent effects on the growth of *P.aeruginosa*. Two different concentrations of GDE, 4mM and 8 mM were tested which are three times and six times higher than the IC_50_ of GDE against the PaSuhB IMPase activity. GDE was not found to impact growth of *P. aeruginosa* significantly at 4mM concentration while at 8mM concentration, GDE inhibits the bacterial growth completely (Figure 7A). Since at 4mM concentration GDE was not impacting the growth of *P. aeruginosa*, this concentration was chosen to further test the impact on the swimming motility and cold sensitivity of *P. aeruginosa*. Intriguingly, 29% reduction in the swimming motility of *P. aeruginosa* was observed upon 4mM GDE treatment (Figure 7B). Moreover, the impact of GDE on cold-sensitive growth was also observed as indicated in Figure 7C1 and 7C2 for *P. aeruginosa* and *E.coli*, respectively. From the figures it is evedent that GDE treated bacteria grow well at 37°C while very little growth was observed at 25°C in *P. aeruginosa* and no growth was observed in case of *E.coli*. The cold sensitive growth pattern was only observed when the bacteria are treated with GDE, these observations are in line with the phenotypes observed with SuhB deletion mutants earlier reported in *E.coli*. These *in-vitro* experiments suggest that GDE binding to PaSuhB may inhibit its regulatory function as extragenic supressor which may further modulate the pathogenesis and some virulence determining factors of *P. aeruginosa*.

**Figure 7:**
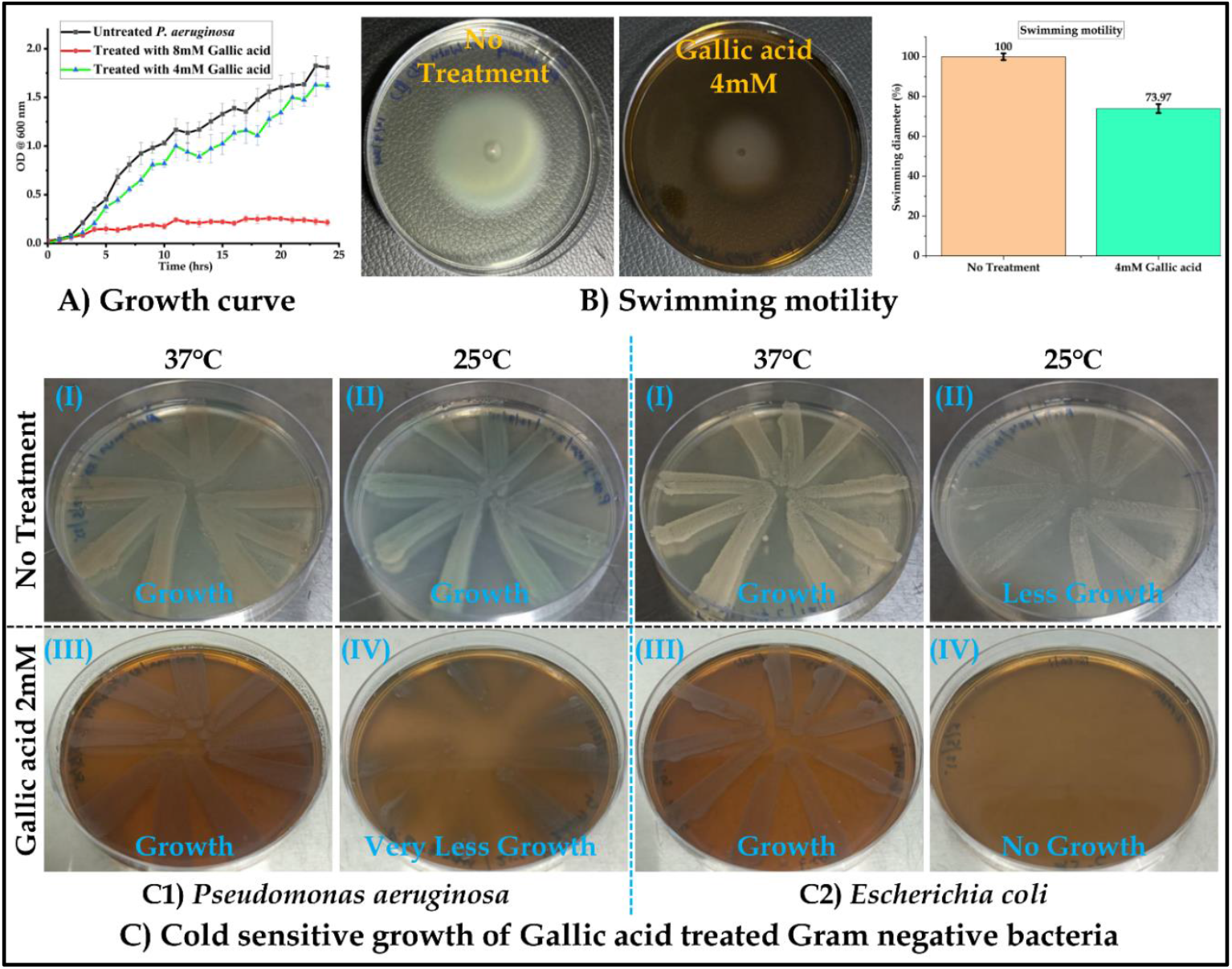
Effect of gallic acid on growth, virulence factor and cold sensitivity of *P. aeruginosa*. Gallic acid effect on A) Growth curve, B) Swimming motility and C) Cold sensitive growth.

## Discussion

*P. aeruginosa*, a Gram-negative opportunistic pathogen from ESKAPE group of bacteria has recently caught global attention due to its emerging roles in myriad reported cases of nosocomial drug-resistant infections with high rates of mortality and morbidities [37-38]. The extensive capabilities of this bacterium to produce numerous virulence factors and the formation of biofilms on biotic/abiotic surfaces play the most crucial role in its pathogenesis and drug resistance [40-42]. The bacterium is also capable to switch from planktonic to biofilm-forming states in response to the appropriate environmental cues. PaSuhB is one of the identified master regulators of *P. aeruginosa* which has been found to exert pleiotropic virulence regulatory function of the organism, ranging from biofilm formation, type -III secretion system, bacterial motility, and antibiotic resistance [25-26]. In other Gram-negative organisms SuhB orthologs have been also found to regulate biofilm formation, exopolysaccharide biosynthesis, r-RNA synthesis and maturation etc. Owing to this pleotropic virulence regulatory function of SuhB protein orthologs, PaSuhB may be considered as one of the lucrative drug target proteins for multiple drug-resistant *P. aeruginosa* strains, amenable to small-molecule-based inhibition. In this present work, we have determined high resolution crystal structures of PaSuhB protein in its apo- and substrate (D-*myo*-inositol 1-monophosphate) bound forms. The crystal structure of PaSuhB was found to be highly reminiscent to the previously reported SuhB and inositol monophosphatase structures with stringent substrate specificity (like E. *coli* SuhB and human IMPases but unlike IMPases orthologues from high-temperature-dwelling bacteria and archaea). Just like its previously identified orthologues, PaSuhB also shows Mg^2+^ activated IMPase activity *in vitro* (Supplementary Figure 2). However, unlike human and other hyper-thermophilic bacterial IMPases, PaSuhB possesses a plausible NusA AR2 domain binding surface just like its *E. coli* counterpart (∼65% amino acid sequence identity). Recently the *E. coli* SuhB dimer has been shown to play an integral role in rRNA transcription anti-termination complex by virtue of its interaction with Nus factors and transcribing RNA polymerase. In this context, the oligomerization state of *E. coli* SuhB was found to be extremely crucial as it governs the interaction with RNA polymerases [22-24] as well as its central interaction in rRNA transcription anti-termination complex. Therefore, any modulation of the oligomeric state of SuhB protein orthologues might result in complete jeopardy in their Nus-dependent pleotropic regulatory functions. Herein we used the high resolution (2.2Å) crystal structure of PaSuhB/substrate binary complex to extract 3D-pharmacophoric features to scout potential small-molecule-based inhibitor leads. Eventually, we identified, GDE, a phyto-phenol naturally abundant in numerous medicinal plants, as one of the most promising inhibitors leads against PaSuhB protein. Biochemical and Biophysical experiments further suggest GDE can bind PaSuhB at low micromolar range and this binding also inhibits the IMPase activity non-competitively. Furthermore, the high-resolution (2.2Å) crystal structure of PaSuhB/GDE complex also corroborates the visual glimpse of two distinct binding sites of GDE inside the PaSuhB active site groove; one at the active site catalytic cleft, near the highly conserved metal binding site and the other at the close proximity of the α4 helix and its preceding loop, distant from the metal binding catalytic cleft. To our knowledge, other than Li^+^ and its structural mimetics which bind at the metal binding catalytic groove of IMPase (SuhB), the discovery of a small molecule inhibitor targeting the active site distal pocket of SuhB (IMPase) protein is novel. Intriguingly, the novel GDE binding site of PaSuhB is also in close proximity to the conserved Arg188 residue (Figure 4 B2, B3 and F2, F3.), which has been previously found to impact oligomerization state of *E. coli* SuhB [24] and thereby can regulate the extragenic suppressor activity of the protein. Importantly, unlike the apo- and substrate bound crystal structures of PaSuhB protein, the asymmetric unit of PaSuhB/GDE complex crystal has an asymmetric tetramer composed of two PaSuhB functional dimers. The formation of this asymmetric PaSuhB tetramers leads to complete shielding of the highly conserved NusA-AR2 binding domain of PaSuhB (Figure 5D.). In solution, dynamic light scattering study also shows that GDE binding to PaSuhB, leads to stable tetramer formation (Figure 6E and F). Altogether, the compelling experimental results suggest that the GDE binding to PaSuhB protein, not only inhibits its enzymatic activity (IMPase) but may also impede its pleotropic virulence regulatory role as an extragenic suppressor. Accordingly, the effect of GDE on the planktonic growth, swimming motility and ‘cold-sensitive growth’ behaviour have been tested in *Pseudomonas aeruginosa*. Interestingly, GDE has been found to induce ‘cold sensitive growth’ in *P. aeruginosa* and *E. coli* [*E. coli* SuhB shares 55.09% overall sequence identity and high degree of structural similarity (RMSD: 0.912Å)] which is the characteristic feature of suhB isogenic null mutant of *E. coli*. Furthermore, GDE was also found to inhibit the swimming motility of the bacteria. The *in vitro* effect of GDE on other virulence determining factors of *P. aeruginosa* and its impact on pre-clinical animal infection model is currently under investigation and will be published elsewhere.

## Conclusions

In the current study, we discovered a small molecule-based inhibitor lead/scaffold (gallic acid) against the pleotropic virulence regulator protein SuhB of *P. aeruginosa* (PaSuhB). Moreover, a novel active site distal, dimerization interface proximal ligand binding site of PaSuhB has been deciphered through high resolution X-ray crystallographic analysis. The ligand gallic acid (GDE) inhibits the enzymatic activity of PaSuhB protein non-competitively, however, most importantly, it also promotes higher order oligomerization of the protein which might be non-compatible for its NusA dependent r-RNA antitermination activity. *In vitro* GDE also found to inhibit the swimming motility of *P. aeruginosa* and induce the ‘cold sensitive growth’ behaviour in *P. aeruginosa* and *E. coli*. Altogether, the present study provides a solid structural-biochemical framework to decipher novel small molecule-based inhibitor lead/scaffold against pleotropic virulence factor PaSuhB, which can be explored further to develop new therapeutic leads against the dreaded multiple drug-resistant *P. aeruginosa* infections.

## Methodology

### PaSuhB purification, crystallization and structure solution

The homogeneously purified and supersaturated PaSuhB was subjected for crystallization trials in sitting drop mode using crystal screen and index solution provided by Hampton Research Institute. Following the initial crystallization hits, fine screening was performed in sodium acetate trihydrate solution of varying pH with PEG 3350 in hanging drop mode in 24 well plates. Co-crystallization with IPD was also done in similar manner with addition to the incubation of protein with 5X concentration of the substrate at room temperature for 30 min. X-ray diffraction data of the single protein crystal in apo form and with substrate was obtained by mounting the single protein crystals (20% glycerol as cryo-protectant) at synchrotron X-ray radiation source (at Indus 2 synchrotron radiation source, RRCAT, Indore, India). Data was collected for 180 degrees with 1 degree rotation for each data collection frame, the detector distance was 260 mm and crystals were subjected to 0.97Å X-ray beam. After collecting the full-length data, the processing was done with different software. Initially the data was reduced and indexed by using XDS software [43] and iMosflm [44]. The reduced data then was subjected to scaling and merging by the pointless and Scala from the CCP4 package [45]. The content of the crystal asymmetric unit was determined by calculation of the Matthews coefficient [46]. Since our protein shares amino acid sequence identity 55.09% with *E. coli* SuhB hence molecular replacement phasing technique was attempted to obtain the molecular densities from the diffraction pattern data using Phaser and MolRep. After getting the initial phase information from molecular replacement phase improvement through iterative cycles of model building and structure refinement was carried out by Coot [47] and Refmac5 and Phenix [48-49] respectively until a good agreement between the experimental data and the structural model is obtained. Finally, the molecular model was subjected to structural validation before submission to protein data bank (PDB; https://www.rcsb.org/). The crystal structure of the apo PaSuhB (PDB ID: 8WIP) and bound with IPD (PDB ID: 8WDQ) was submitted to PDB.

### *In-silico* pharmacophore guided similarity search for plausible inhibitors against PaSuhB active site

The concept of structure [50]and pharmacophore [51] guided drug discovery was employed to screen plausible inhibitor against our target protein PaSuhB. PaSuhB substrate bound structure provides glimpse of interaction types at its active site which further exploited for inhibitor search. Here, the pharmacophore information of D-*myo*-Inositol 1-monophosphate (PaSuhB substrate) was harnessed to search the library of similar compounds using SwissSimilarity server against bioactive molecules. This *in-silico* screening results hundreds of similar molecules with polyphenol/polyhydroxy phosphate and carboxylate moiety. Since, phosphates are the substrate of this enzyme and nonspecific phosphatases abundantly present in physiological conditions, we have selected only molecules with polyphenol/polyhydroxy carboxylates for further *in-vitro* inhibition studies.

### PaSuhB IMPase activity inhibition with polyphenol and poly hydroxy carboxylate compounds

Purified PaSuhB hydrolyses IPD, hence to test the enzyme activity inhibition we have utilized IPD as substrate and malachite green phosphate assay kit for *in-vitro* screening of inhibitors. We have used enzyme 0.25µM, Mg^2+^ 30mM, substrates 100 µM and reaction was carried out in buffer containing 20 mM tris pH 8.0 with 150 mM NaCl and varying concentration of inhibitors, reaction mixture was incubated at 37°C for 2 min. After the completion of reaction, we added 50 µl of malachite green solution containing 100 parts of reagent A and 1 part of reagent B followed by incubation at room temperature for 30 min and absorbance measurement at 620 nm. We also performed the mode of inhibition study with the most potent inhibitor GDE (2.5mM) with varying substrate concentrations.

### Fluorescence based protein ligand interaction study

To analyse the ligand protein interaction behaviour and investigation of number of binding sites we exploited the quenching of intrinsic fluorescence properties of PaSuhB. Since PaSuhB have several aromatic amino acids including tryptophan it gives strong fluorescence spectra at 310-350nm. For the fluorescence assay to probe PaSuhB interaction with our ligand (GDE), the purified PaSuhB (0.5µM) titration assay was performed with varying concentration of GDE using spectrofluorometer (JASCO FP8300). The reaction mixtures were incubated at room temperature for 20 minutes and then the fluorescence readings were taken. The excitation and emission band width were fixed to 10nm, the scan speed was 200nm min^-1^. The fluorescence emission spectra were recorded from 300 nm to 600 nm while the excitation was fixed at 280 nm. The results of fluorescence spectra were plotted using origin software [52] and GraphPad Prism [53].

#### PaSuhB and GDE co-crystallization and structure solution

From the biochemical experiments it was evident that GDE interacts with PaSuhB and inhibits its enzymatic activity at sub millimolar concentration. Next, we moved towards crystallization of PaSuhB in complex with GDE. Since crystals of PaSuhB apo form and with substrates appears in 0.1M sodium acetate trihydrate pH 5.0 and 4.5 respectively with PEG3350 20%, so we have tried these conditions to crystallize PaSuhB in complex with GDE also. The concentrated protein (∼20 mg/ml) was mixed gently with 10X concentration of GDE and incubated for 30 min at room temperature. Further, equal volume of the crystallization solution and protein-ligand mix then subjected for crystallization in hanging drop mode in 24 well plates. The diffraction quality crystals were appeared after 2 days of incubation in 0.1M sodium acetate pH 5.0 and 20% PEG3350. X-ray diffraction of crystals, data collection, and structure solution was done in similar manner as apo form and IPD complex with minor modifications. The 10% glycerol was used as cry protectant; detector distance was kept 180 mm for 2.2Å and 200 mm for 2.8Å resolution crystals. Molecular replacement was done with apo from of PaSuhB crystal, the molecular model was subjected to structural validation before submission to protein data bank (PDB; https://www.rcsb.org/). Two crystal structures of PaSuhB bound with GDE was submitted to PDB (PDB: 9JFC and 9J90) with 2.2 and 2.8Å resolution respectively.

### Molecular dynamic simulation study and hydrodynamic diameter measurement

To investigate the dynamics of PaSuhB apo structure and PaSuhB-GDE complexes in physiologically simulated conditions molecular dynamic (MD) simulation was performed using NAMD for the time scale of 30 ns [54]. Parameter and topology files of ligands and proteins were generated using CHARM-GUI [55] and Visual molecular dynamic (VMD) [56] respectively. To solvate the system, default protein dimensions was applied to cubic water cells and to set isothermal-isobaric ensemble environment, Langevin dynamic was applied. Energy minimization was done according to steepest descent method for 1000 steps; non-bonded interaction was calculated within a cut-off of 10Å and with 3000 steps of frequency for restart and dcd. The time for each step was set 2 femtoseconds throughout whole simulation process and electrostatic interaction was calculated using Particle Mess Ewald (PME) method. Post completion of simulation, from the generated trajectories RMSD, RMSF, Rg and SASA were analysed using VMD software. To investigate the oligomerization status of PaSuhB upon GDE treatment the hydrodynamic diameter was measured using Zetasizer instrument (Malvern Panalytical). Briefly, 50µM of PaSuhB was incubated in 10mM Tris pH 8.0 and 100mM NaCl with 8mM of GDE for 30 min at room temperature followed by size measurement. The size of untreated PaSuhB was measured as control to visualize the impact of GDE on in solution oligomerization status of PaSuhB.

### Growth curve, swimming motility and cold sensitivity assay

LB broth 200 µl was inoculated with 1% overnight culture of *P. aeruginosa* in 96 well plates with 4mM gallic acid (pH 7.4) treatment, no treatment was kept as control. All experiments were carried out in triplicates. The plates were incubated in photometer for 24 hrs at 37 °C with 150 rpm shaking. Optical density (OD) at 600 nm was measured at each hour of interval till 24 hrs. Graph of growth curve was plotted against OD versus time using OriginPro 2025. In order to test the bacterial swimming motility [57], we have inoculated the 5 µl of 5 times diluted overnight grown *Pseudomonas aeruginosa* PAO1 culture into center of the 0.3% agar prepared in LB broth. To see the effects of 4mM GDE along with no treatment control where water added in compensation of the ligands volume. All the ligands were dissolved in milli-Q water followed by sterilization with 0.22µ filter paper. The plates were incubated at 37°C for 16 hours followed by measurement of the swim zone diameter and results analysis. Cold sensitive growth study [58] was done on 2% LB agar plates with 2, 3 and 4mM of Gallic acid treatment along with no treatment control. Log phase *P. aeruginosa* culture was streaked on agar plates followed by incubation at 37°C and 25 °C separately. After 24 hours of incubation plates were observed and photos of the plates are taken with personal cell phone camera.

## Data availability

Structure coordinates and map files are deposited into worldwide protein data bank (wwPDB) with the following accession codes, PDB IDs 8WIP, 8WDQ, 9JFC and 9J90. Source data are provided in this paper and supplementary information. Raw experimental data and analyzed reports are available with corresponding author which can be access upon request as per journal policy.

## Acknowledgements

We thank IIT Jodhpur for providing infrastructures and experimental facility to carryout experiments cited in this manuscript. AYURTECH facility at IIT Jodhpur, Ministry of AYUSH, Government of India for providing financial assistance (Sanction number: S-12011/12/2021-SCHEME). RRCAT Indore, India for providing Synchrotron radiation source for protein crystal diffraction and data collection. DMD facility at IIT Jodhpur, funded by SERB, Department of Science and Technology, Government of India. Authors including V.K.Y and A.J express special thanks to Ministry of Education, Government of India for providing fellowships to carryout doctoral research. We acknowledge the Microsoft Edge based Copilot AI tool for taking assistance for editing and formatting of this manuscript.

## Author’s contributions

This project was conceived and designed by S.B, V.K.Y and M.M. V.K.Y has conducted all the experiments unless otherwise mentioned specifically, analysed data, prepared figures, wrote complete manuscript and finalized that. Molecular dynamic simulation study was conducted, analysed and explained by A.J along with that they also assisted in protein crystallization and data collection. The manuscript is edited and finalized by S.B and M.M.

## Additional information

Additional informations are available in supplementary data

## References

1. Wilson, M. G. and Pandey, S. Pseudomonas aeruginosa. 2023 Aug 8. In: StatPearls [Internet]. Treasure Island (FL): StatPearls Publishing; 2025 Jan–. PMID: 32491763.

2. Urgancı, N. N. et al. Pseudomonas aeruginosa and its pathogenicity. Turkish Journal of Agriculture-Food Science and Technology 10(4), 726–38 (2022).

3. Ali, N. M. et al. Pseudomonas aeruginosa-associated acute and chronic pulmonary infections. Pathogenic Bacteria. IntechOpen, (2020).

4. Turner, K. H. et al. Requirements for Pseudomonas aeruginosa acute burn and chronic surgical wound infection. PLoS genetics 10(7), e1004518 (2014).

5. Glen, K. A. et al. β-lactam Resistance in Pseudomonas aeruginosa: Current Status, Future Prospects. Pathogens 10(12), 1638 (2021).

6. Horcajada, J. P. et al. Epidemiology and treatment of multidrug-resistant and extensively drug-resistant Pseudomonas aeruginosa infections. Clinical microbiology reviews 32(4), 10–128 (2019).

7. https://www.who.int/news/item/27-02-2017-who-publishes-list-of-bacteria-for-which-new-antibiotics-are-urgently-needed.

8. Irvine, R. F. et al. Inositol phosphates: proliferation, metabolism and function. Philosophical Transactions of the Royal Society of London. B, Biological Sciences 320(1199), 281–98 (1988).

9. Kim, S. et al. The inositol phosphate signalling network in physiology and disease. Trends in Biochemical Sciences 49(11), 969–85 (2024).

10. Roberts, M. F. Inositol in bacteria and archaea. Biology of Inositols and Phosphoinositides: Subcellular Biochemistry 1, 103–33 (2006).

11. Matsuhisa, A. et al. Inositol monophosphatase activity from the Escherichia coli suhB gene product. Journal of bacteriology 177(1), 200–5 (1995).

12. Movahedzadeh, F. et al. Inositol monophosphate phosphatase genes of Mycobacterium tuberculosis. BMC microbiology 10, 1–5 (2010).

13. Bhattacharyya, S. et al. Crystal structure of Staphylococcal dual specific inositol monophosphatase/NADP (H) phosphatase (SAS2203) delineates the molecular basis of substrate specificity. Biochimie 94(3), 879–90 (2012).

14. Bhattacharyya, S. et al. Structural elucidation of the NADP (H) phosphatase activity of staphylococcal dual-specific IMPase/NADP (H) phosphatase. Biological Crystallography 72(2), 281–90, (2016).

15. Pirovich, D. B. et al. Multifunctional fructose 1, 6-bisphosphate aldolase as a therapeutic target. Frontiers in molecular biosciences 8, 719678 (2021).

16. Gizak, A. et al. Fructose 1, 6-bisphosphatase as a promising target of anticancer treatment. Advances in Biological Regulation 23, 101057 (2024).

17. Can, A. et al. Molecular actions and clinical pharmacogenetics of lithium therapy. Pharmacology, biochemistry, and behavior 123, 3–16 (2014).

18. Rosales, R. R. et al. The suhB gene of Burkholderia cenocepacia is required for protein secretion, biofilm formation, motility and polymyxin B resistance. Microbiology 158(9), 2315–24 (2012).

19. Boles, B. R. et al. Identification of genes involved in polysaccharide-independent Staphylococcus aureus biofilm formation. PloS one 5(4), e10146 (2010).

20. Janczarek, M. and Skorupska, A. Regulation of pssA and pssB gene expression in Rhizobium leguminosarum bv. trifolii in response to environmental factors. Antonie Van Leeuwenhoek 85(3), 217–27 (2004).

21. Movahedzadeh, F. et al. Inositol monophosphate phosphatase genes of Mycobacterium tuberculosis. BMC microbiology 10(1) 1–15 (2010).

22. Huang YH, et al. Structure-based mechanisms of a molecular RNA polymerase/chaperone machine required for ribosome biosynthesis. Molecular cell 79(6), 1024–36 (2020).

23. Huang, Y. H. et al. Structural basis for the function of SuhB as a transcription factor in ribosomal RNA synthesis. Nucleic acids research 47(12), 6488–503 (2019).

24. Wang, Y. et al. The structure of the R184A mutant of the inositol monophosphatase encoded by suhB and implications for its functional interactions in Escherichia coli. Journal of Biological Chemistry 282(37), 26989–96 (2007).

25. Li, K. et al. SuhB is a regulator of multiple virulence genes and essential for pathogenesis of Pseudomonas aeruginosa. MBio 4(6), e00419–13 (2013).

26. Li, K. et al. SuhB regulates the motile-sessile switch in Pseudomonas aeruginosa throughthe Gac/Rsm pathway and c-di-GMP signaling. Frontiers in microbiology 8, 1045 (2017).

27. Shi, J. et al. SuhB is a novel ribosome associated protein that regulates expression of MexXY by modulating ribosome stalling in Pseudomonas aeruginosa. Mol Microbiol 98(2), 370–83 (2015).

28. Shamsi, S. et al. Stability, Toxicity, and Antibacterial Potential of Gallic Acid-Loaded Graphene Oxide (GAGO) Against Methicillin-Resistant Staphylococcus aureus (MRSA) Strains. Int J Nanomedicine 17, 5781–5807 (2022).

29. Abdella, M. et al. Antibacterial Evaluation of Gallic Acid and its Derivatives against a Panel of Multi drug resistant Bacteria Med Chem 20(2), 130–139 (2024).

30. Chaiprateep, E O., et al. Formulation of triphala-based products: Focus on gallic acid content and health benefits. J Adv Pharm Technol Res 16(4), 213–218 (2025).

31. Phansawan, B. et al. Determination of gallic acid and rutin in extracts Cassia alata and Andrographis paniculata.” Science Asia 40 414–419 (2014).

32. Kahkeshani, N. et al. Pharmacological effects of gallic acid in health and diseases: A mechanistic review. Iran J Basic Med Sci. 22(3), 225–237 (2019).

33. Keyvani, G. S. et al. An update on the potential mechanism of gallic acid as an antibacterial and anticancer agent. Food Science & Nutrition 11(10), 5856–72 (2023).

34. Tian, Q. et al. Bactericidal activity of gallic acid against multi-drug resistance Escherichia coli. Microbial Pathogenesis 173, 105824 (2022).

35. Zoete, V. et al. SwissSimilarity: a web tool for low to ultrahigh throughput ligand-based virtual screening. Journal of Chemical Information and Modeling 56(8), 1399–1404 (2016).

36. Koes, D. R. and Camacho, C. J. ZINCPharmer: pharmacophore search of the ZINC database. Nucleic acids research 40(W1), W409–14 (2012).

37. Stieglitz, K. A. et al. Crystal structure of a dual activity IMPase/FBPase (AF2372) from Archaeoglobus fulgidus: The story of a mobile loop. Journal of Biological Chemistry 277(25), 22863–74 (2002).

38. Brown, A.K. et al. Dimerization of inositol monophosphatase Mycobacterium tuberculosis SuhB is not constitutive, but induced by binding of the activator Mg^2+^. BMC Struct Biol 7, 55 (2007).

39. Narimisa, N. et al. Prevalence of colistin resistance in clinical isolates of Pseudomonas aeruginosa: a systematic review and meta-analysis. Front Microbiol 15, 1477836 (2024).

40. Valzano, F. et al. Resistance to ceftazidime–avibactam and other new β-lactams in Pseudomonas aeruginosa clinical isolates: a multi-center surveillance study. Microbiology Spectrum 12(8), e04266–23. (2024).

41. Liao, C. et al. Virulence factors of Pseudomonas aeruginosa and antivirulence strategies to combat its drug resistance. Frontiers in cellular and infection microbiology 12, 926758 (2022).

42. Thi, M.T. et al. Pseudomonas aeruginosa biofilms. International journal of molecular sciences 21(22), 8671 (2020).

43. Kabsch, W. xds. Biological crystallography 66(2), 125–32 (2010).

44. Kabsch, W. Integration, scaling, space-group assignment and post-refinement. Acta Crystallographica Section D: Biological Crystallography 66(2), 133–144 (2010).

45. Project, Collaborative Computational. The CCP4 suite: programs for protein crystallography. Acta crystallographica. Section D, Biological crystallography 50(5), 760–763 (1994).

46. Kantardjieff, K. A. and Bernhard R. Matthews coefficient probabilities: improved estimates for unit cell contents of proteins, DNA, and protein–nucleic acid complex crystals. Protein Science 12(9) 1865–1871 (2003).

47. Emsley, P. and Kevin, C. Coot: model-building tools for molecular graphics. Acta crystallographica section D: biological crystallography 60(12), 2126–2132 (2004).

48. Nicholls, R. A. et al. Low-resolution refinement tools in REFMAC5. Acta Crystallographica Section D: Biological Crystallography 68(4), 404–417 (2012).

49. Terwilliger, T. C. et al. Macromolecular structure determination using X-rays, neutrons and electrons: recent developments in. Acta Crystallographica Section D Structural Biology 75(10), 861–77 (2019).

50. Batool, M. et al. Structure-Based Drug Discovery Paradigm. Int J Mol Sci. 20(11), 2783 (2019).

51. Gao, Q. et al. Pharmacophore based drug design approach as a practical process in drug discovery. Curr Comput Aided Drug Des. 6(1), 37–49 (2010).

52. Origin, Version 2022. OriginLab Corporation, Northampton, MA, USA.

53. GraphPad Software, Boston, Massachusetts USA, https://www.graphpad.com.

54. Phillips, J C., et al. Scalable molecular dynamics on CPU and GPU architectures with NAMD.” The Journal of chemical physics 153(4) 044130 (2020).

55. Jo, S. K. et al. CHARMM-GUI: a web-based graphical user interface for CHARMM. Journal of computational chemistry 29(11), 1859–1865 (2008).

56. Humphrey, W. et al. VMD: visual molecular dynamics. Journal of molecular graphics 14(1), 33–38 (1996).

57. Khayat, M. T. et al. Sodium citrate alleviates virulence in Pseudomonas aeruginosa. Microorganisms 10(5), 1046 (2022).

58. Chen, L. and Roberts, M. F. Overexpression, purification, and analysis of complementation behavior of E. coli SuhB protein: comparison with bacterial and archaeal inositol monophosphatases. Biochemistry 39(14), 4145–53 (2000).

