## Supplementary Information for "Suppressing the suppressor: Gallic acid induced asymmetric tetramerization of the pleotropic virulence factor SuhB from *Pseudomonas aeruginosa* abolishes its extragenic suppressor activities. A structure-based functional study"

### Table of Contents


| S. No. | Contents | Initial page |
|---|---|---|
| 1 | Figure S1 | 2 |
| 2 | Figure S2 | 2 |
| 3 | Figure S3 | 3 |
| 4 | Tabl2 S1 | 3 |
| 5 | Figure S4 | 4 |
| 6 | Figure S5 | 4 |
| 7 | Figure S6 | 5 |
| 8 | Figure S7 | 5 |
| 9 | Figure S8 | 6 |
| 10 | Figure S9 | 7 |
| 11 | Figure S10 | 8 |
| 12 | Figure S11 | 9 |

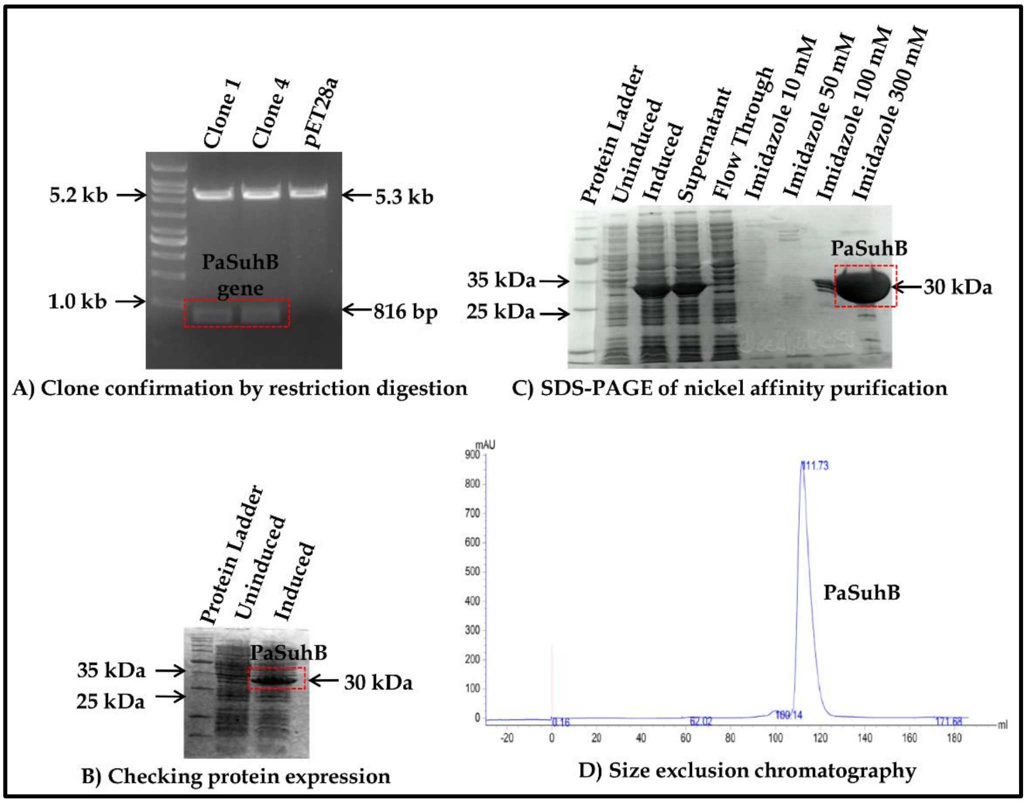


**Figure S1:** PaSuhB cloning, overexpression and purification, A) Represent PaSuhB clone confirmation into pET28a vector through restriction digestion with *Bam*HI and *Hin*DIII enzymes. B) Protein expression checks at 37°C inductions with 100µM IPTG. C) SDS-PAGE analysis of purified PaSuhB fractions through Ni-NTA column chromatography. D) Size exclusion chromatography purification of 300mM imidazole fraction of PaSuhB.

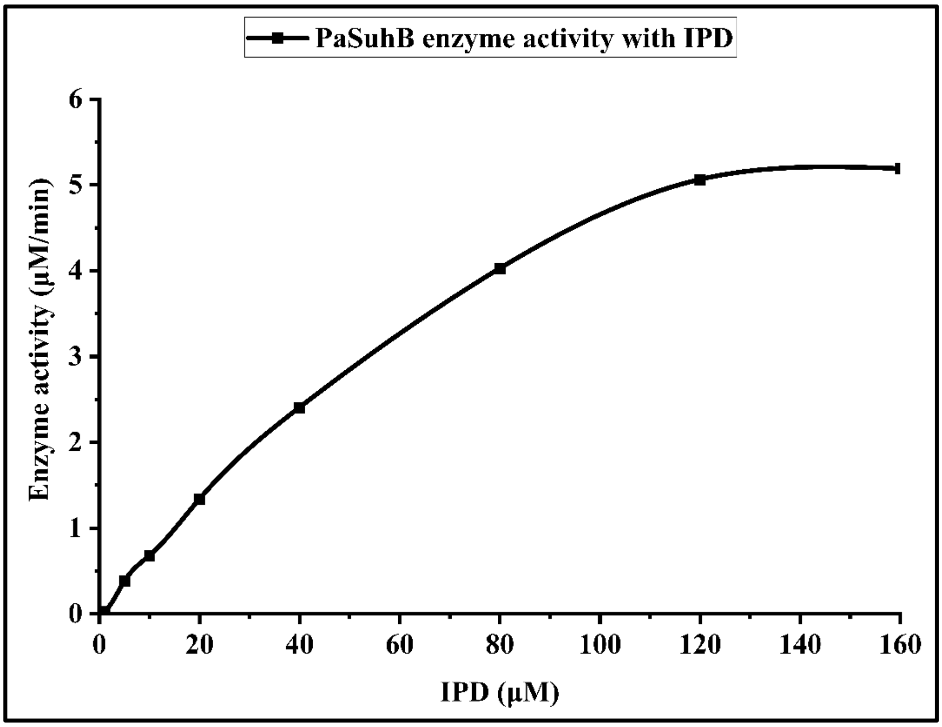


**Figure S2:** $Mg^{2+}$ dependent D-*myo*-inositol 1-monophosphate (IPD) phosphatase activity of purified PaSuhB.

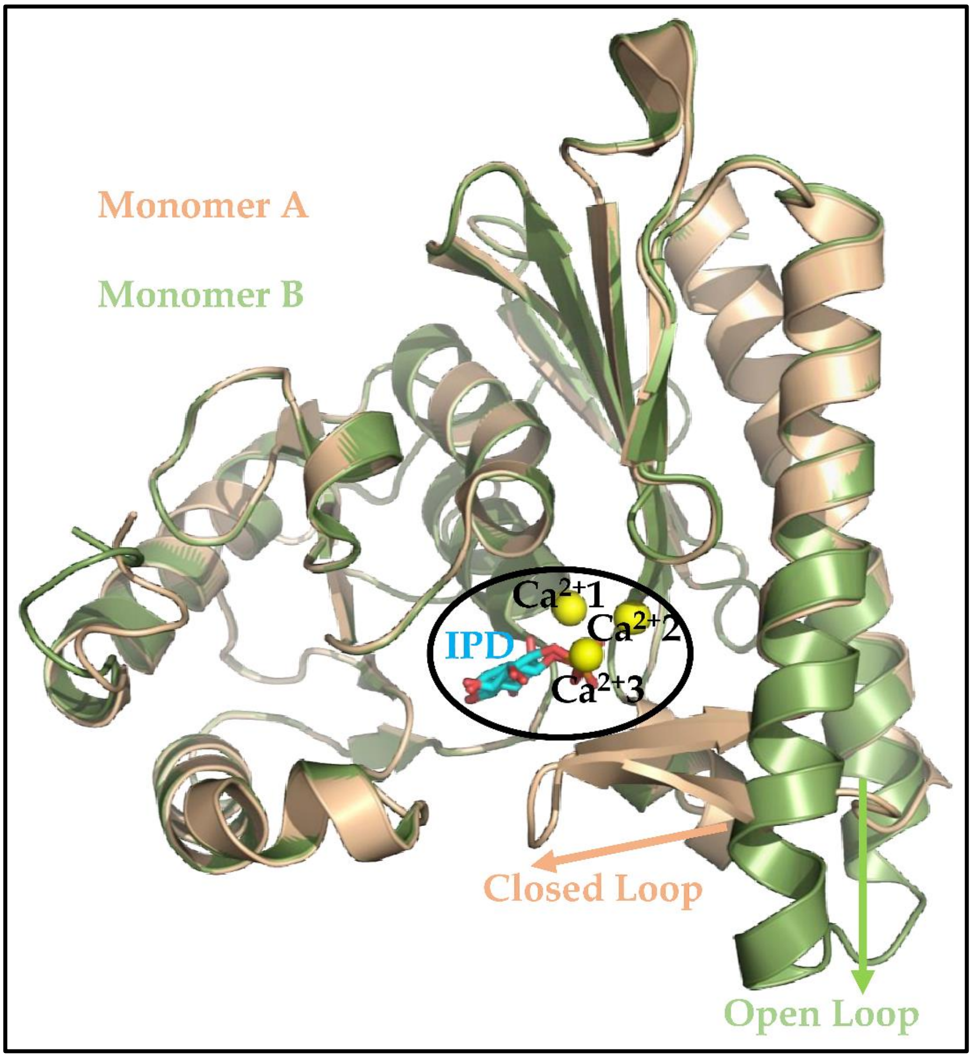


**Figure S3:** Two monomers of PaSuhB-$Ca^{2+}$-IPD complex crystal structure superimposed on each other to indicates closed conformation of active site mobile loop in monomer A (with three metal)

**Table S1: Compounds and their respective $IC_{50}$ value**

| S.No. | Compounds | $IC_{50}$ (mM) | Rank based on $IC_{50}$ |
|---|---|---|---|
| 1 | Alpha ketoglutarate | 59.35 | 7 |
| 2 | Ascorbate | 157 | 14 |
| 3 | Citrate | 20.61 | 3 |
| 4 | Fumarate | 107.5 | 11 |
| 5 | Gallate (GDE) | 1.354 | 1 |
| 6 | Gluconate | 76.35 | 8 |
| 7 | Glucuronate | 118.8 | 13 |
| 8 | Isocitrate | 54.25 | 6 |
| 9 | Malate | 109.3 | 12 |
| 10 | Oxaloacetate | 31.55 | 4 |
| 11 | Pyruvate | 78.56 | 9 |
| 12 | Salicylate | 14.65 | 2 |
| 13 | Shikimate | 41 | 5 |
| 14 | Succinate | 105.5 | 10 |
| 15 | Acetate | No inhibition | 15 |

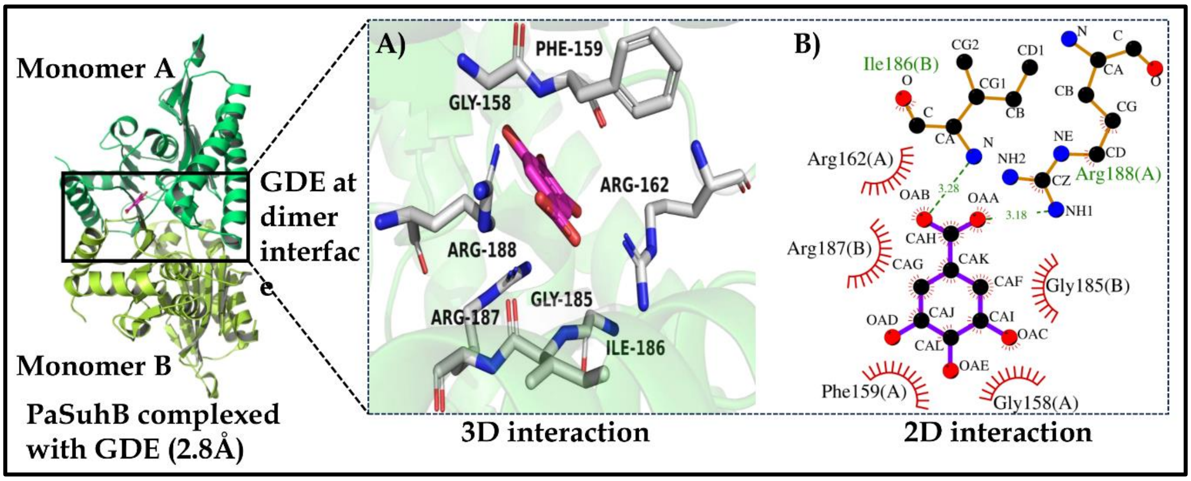


**Figure S4:** Crystal structure of PaSuhB at low resolution (2.8Å) complexed with GDE and their 3D and 2D interaction profiles.

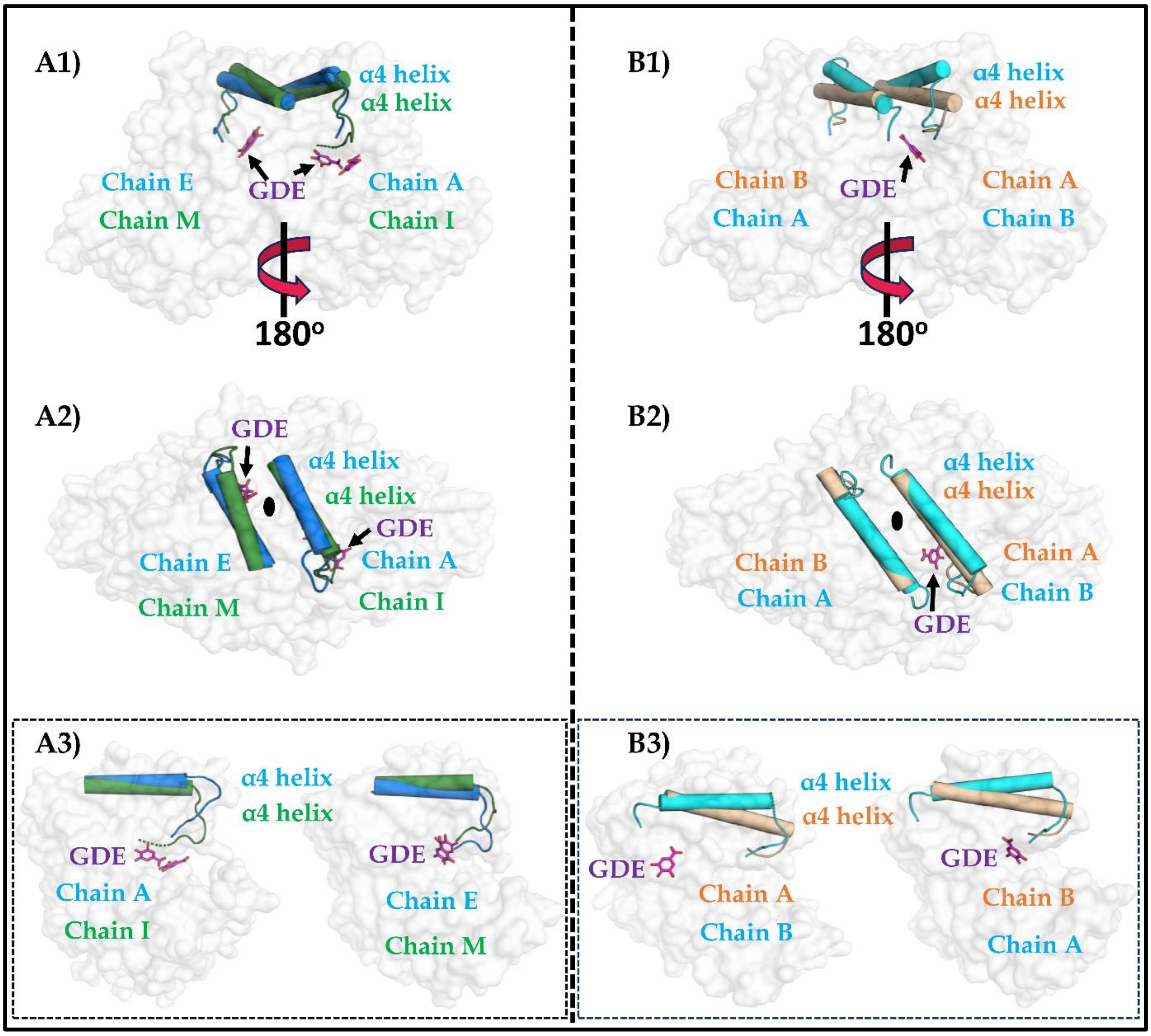


**Figure S5:** GDE induced 3D structural disposition of α4 helix in PaSuhB-GDE complex high resolution (2.2Å) and low resolution (2.8Å) crystal structures. Figure A1-A3 represent comparison of AE dimer with IM dimer of high-resolution structure, A1) Top view, A2) Side view and A3) Respective monomers. Figure B1-B3 represent comparison of AB dimer of PaSuhB apo form with BA dimer of PaSuhB-GDE complex low resolution crystal structure, B1) Top view, B2) Side view and B3) Respective monomers.

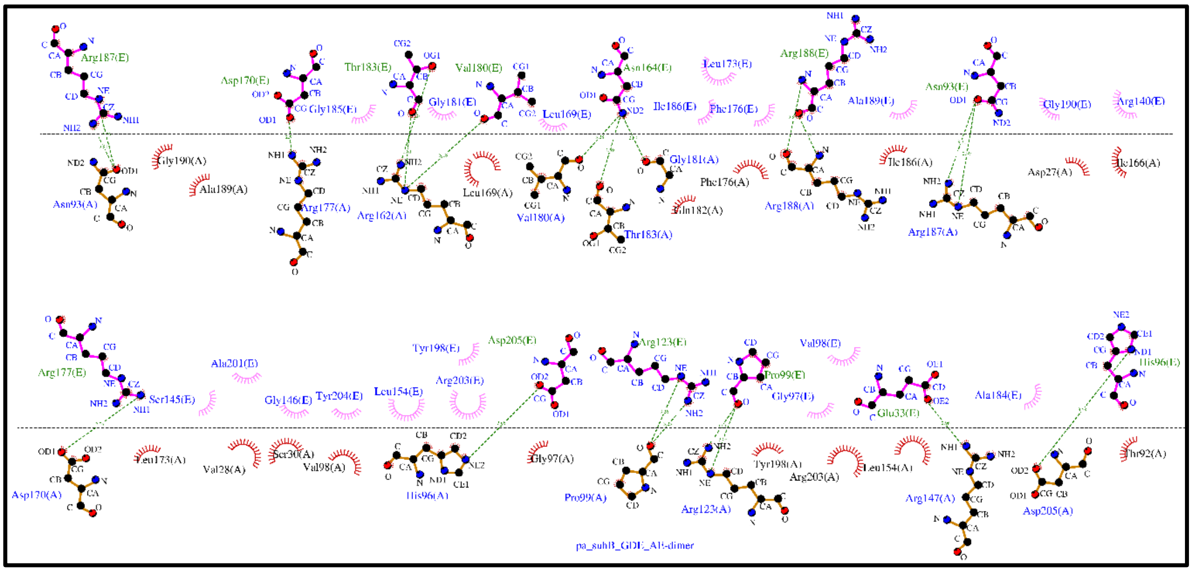

**Figure S6:** Representations of the PaSuhB dimerization residues retrieve from AE dimer of PaSuhB-GDE complex (PDB ID: 9JFC).

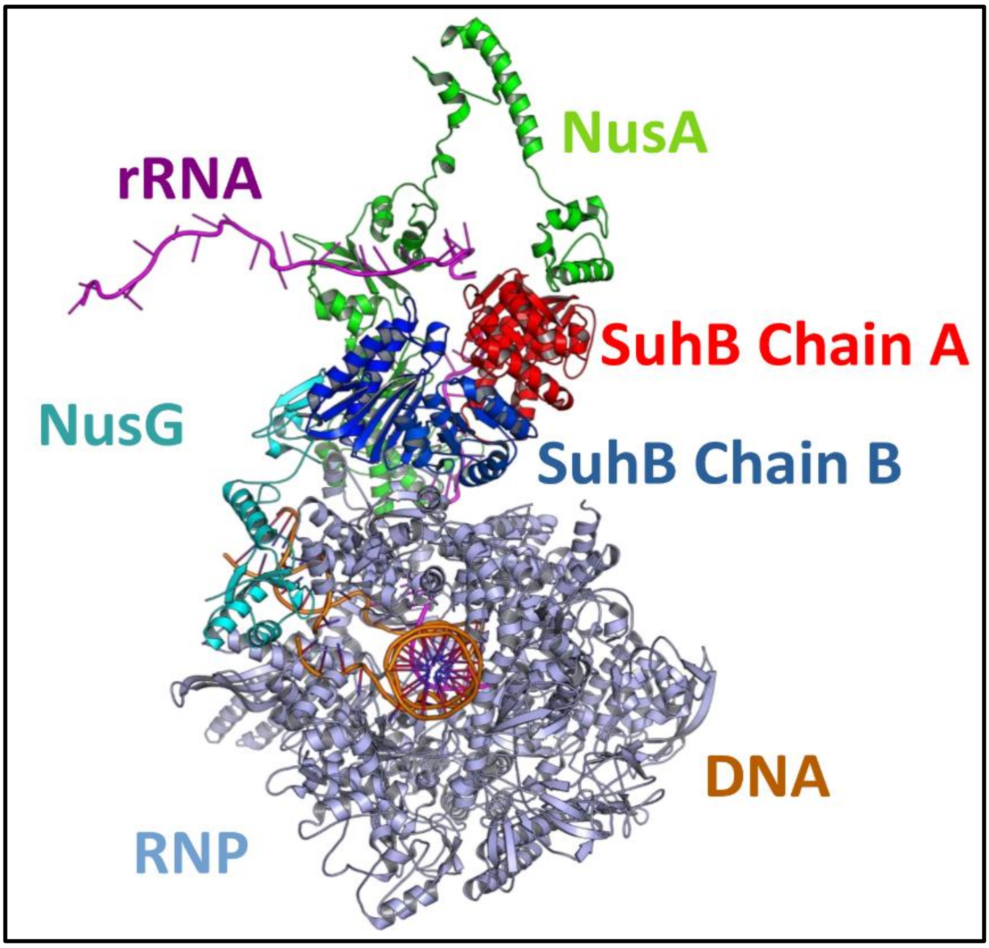


**Figure S7:** Structure of *E. coli* transcription anti-terminator complex presenting SuhB (Red-Blue dimer) interacting partners and their interacting domains.

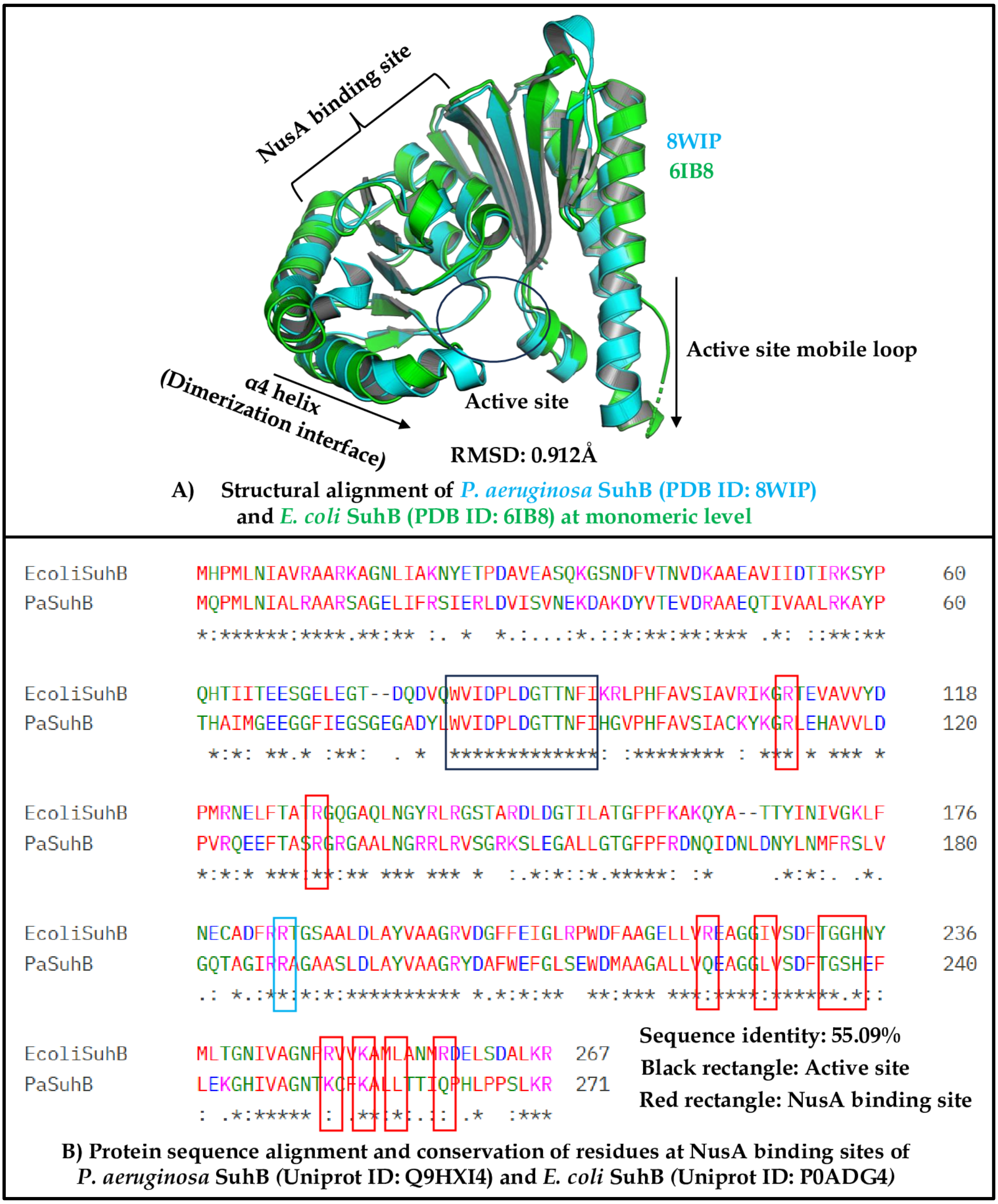


**Figure S8:** Alignment of *P. aeruginosa* SuhB with *E. coli* SuhB based on, A) 3D Structures and B) Amino acid sequences. Cyan rectangle in figure B represent Arginine which is responsible for oligomeric status of SuhB in *E. coli.*

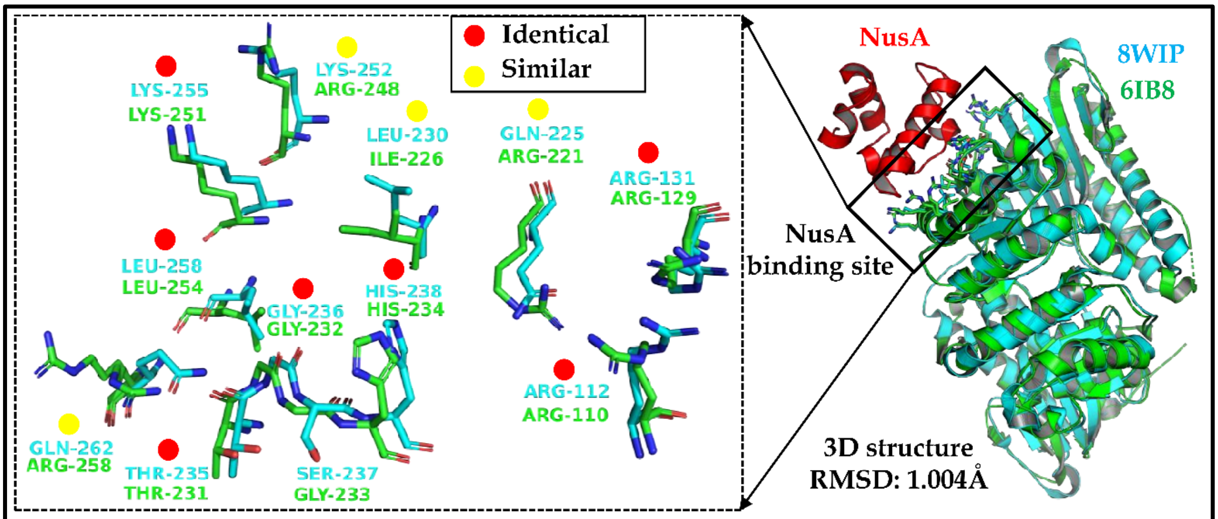


**A) Structural alignment and conservation of residues at NusA binding sites of *P. aeruginosa* SuhB (PDB ID: 8WIP) and *E. coli* SuhB (PDB ID: 6IB8)**

```
Ecoli_NusA_AR2_domain      DNKPADDLLNLEGVDRDLAFKLAARGVCTLEDLAEQGIDDLADIEGLTDEKAGALIMAAR 60
Pa_NusA_AR2_domain         DAQPAEDLLSLDGMSKELALDLALRGVTTREDLAEQSIDDLLDIDGMDEERAGKLIMAAR 60
                           * :**:***.*:*:.::**:.** *** * ******.**** **:*: :*:** ******

Ecoli_NusA_AR2_domain      NICWFGDEA    69
Pa_NusA_AR2_domain         A-HWFE---    65
                             **
```

**Amino acid sequence identity: 65.11%**

**B) Amino acid sequence alignment and conservation of residues at SuhB binding sites of NusA AR2 domain of *P. aeruginosa* (Uniprot ID: Q9HV54) and *E. coli* (Uniprot ID: P0AFF6)**

**Figure S9:** Alignment of *Pseudomonas aeruginosa* with *E. coli*, A) SuhB based on 3D Structures of dimers and B) NusA AR2 domain amino acid sequences.

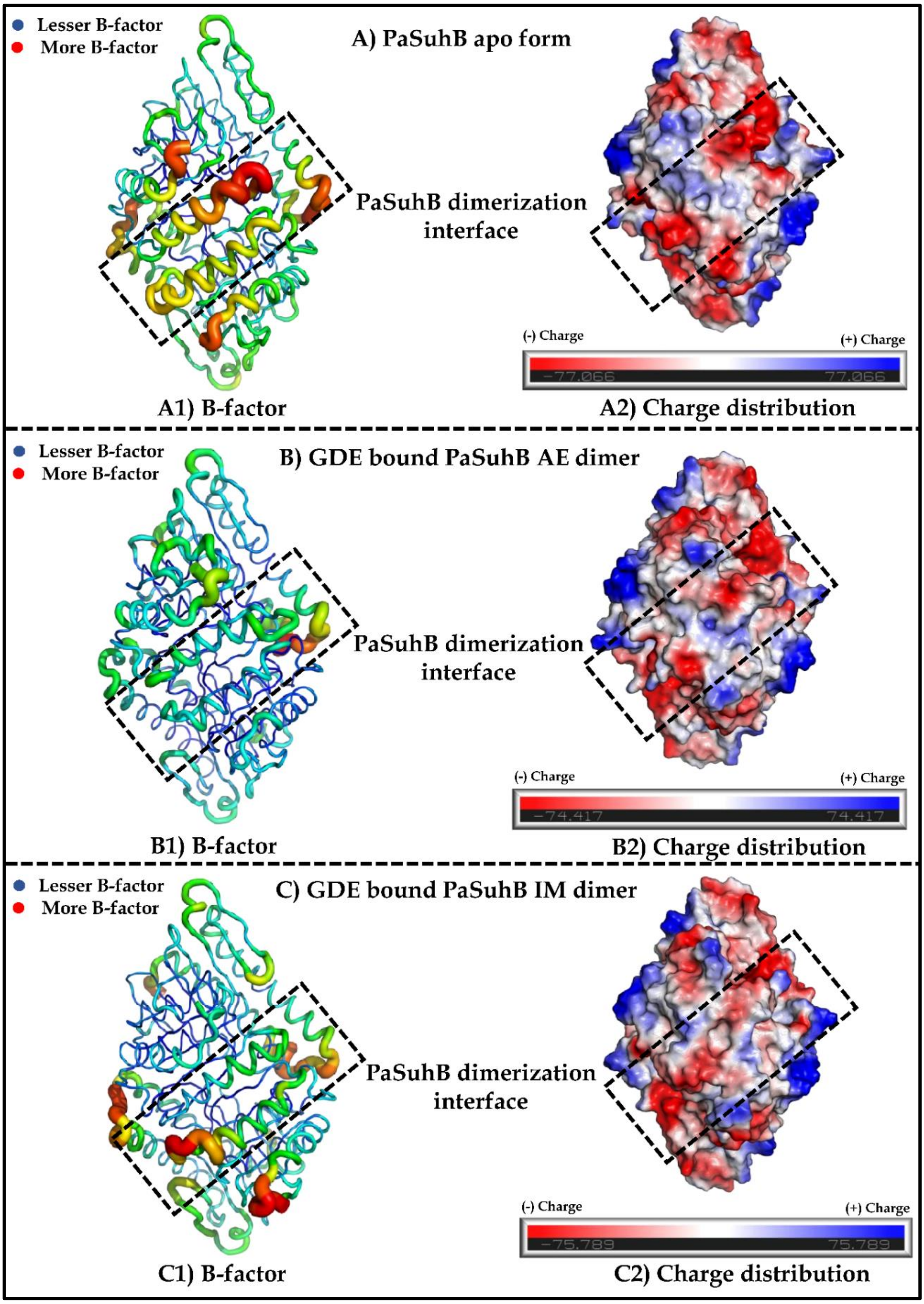


**Figure S10:** Impact of GDE binding on PaSuhB stability and charge distribution of α4 helix, A) and A2) represent temperature factor and charge distribution respectively of PaSuhB apo form crystal structure. Similarly, B1), B2) and C1, C2 represent for GDE bound PaSuhB AE dimer and IM dimer respectively. In figure A1, B1, and C1 blue colour indicates lesser B-factor while red colour indicates higher B-factor, similarly, in figure A2, B2 and C2 blue and red colour represents positive and negative charged amino acid residues respectively.

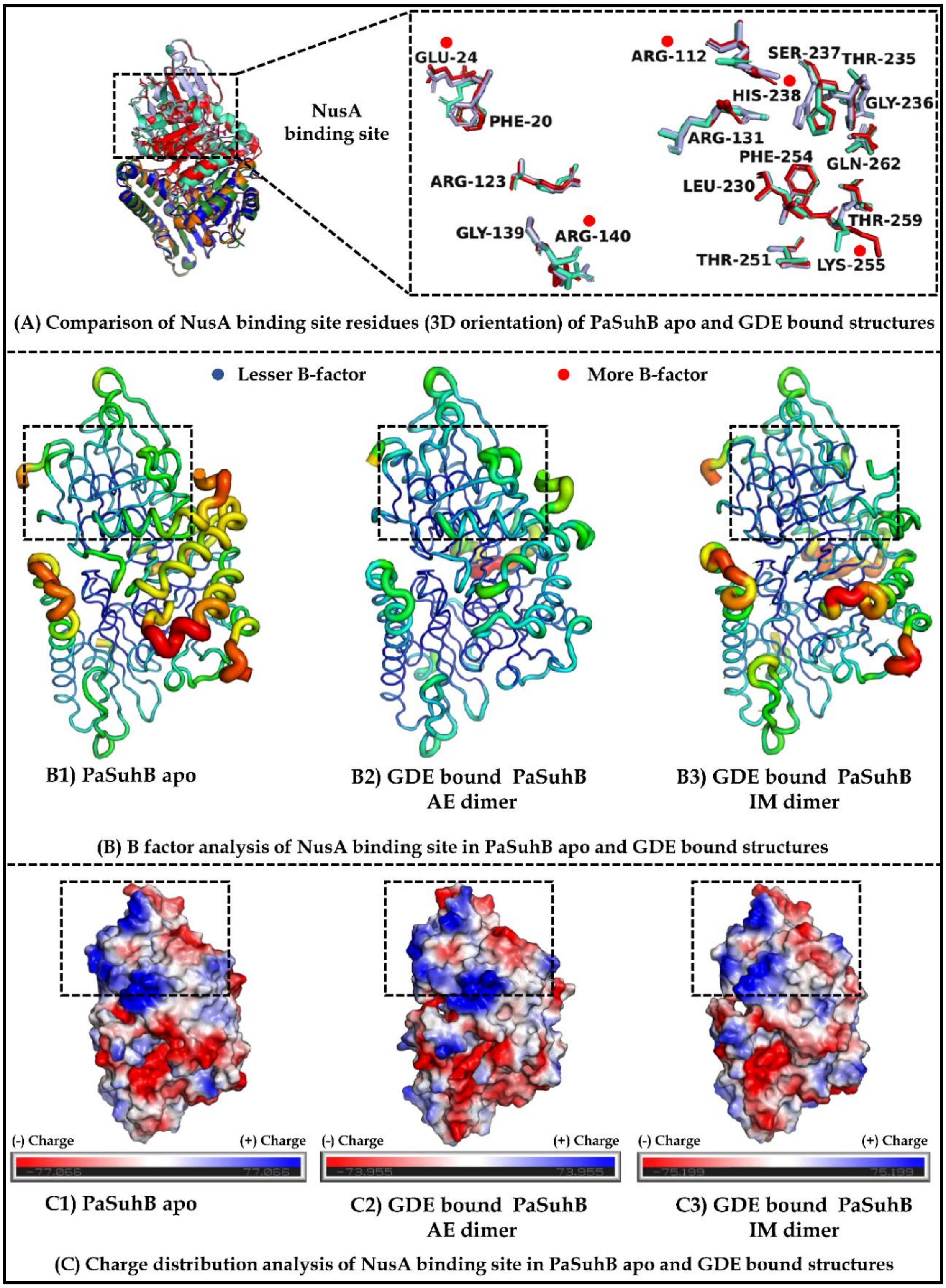


**Figure S11:** Impact of GDE binding on stability and charge distribution of PaSuhB crystal structures at NusA binding site (Tetramerization surface), A1) Represents differential 3D orientation of residues at tetramerization surface of apo (green) and GDE bound PaSuhB dimers (light blue and red), residues indicated with red dots (figure A right panel) show high degree of structural alteration, B) and C) represent stability and charge distribution respectively for AB dimer of PaSuhB apo structure and AE and IM dimers of GDE bound PaSuhB structure. In figure B, blue colour indicates lesser B-factor while red colour indicates higher B-factor, similarly, in figure C, blue and red colour represents positive and negative charged amino acid residues respectively.
